# Bis-hydroxylation of Homocitrulline Catalyzed by a Multinuclear Nonheme Iron-Dependent Oxidative Enzyme during RiPP Biosynthesis

**DOI:** 10.64898/2026.05.01.722236

**Authors:** Dayna P. Hebron, Tucker J. Shriver, Joshua J. Ziarek, Amy C. Rosenzweig

**Author notes:** **Corresponding Author Dayna P. Hebron -** Departments of Molecular Biosciences and of Chemistry, Northwestern University, Evanston, Illinois 60208, United States;, **Amy C. Rosenzweig -** Departments of Molecular Biosciences and of Chemistry, Northwestern University, Evanston, Illinois 60208, United States.

## Abstract

Ribosomally synthesized and post-translationally modified peptides (RiPPs) are produced by biosynthetic enzymes that modify genetically encoded precursor peptide backbones and side chains. Genome mining and bioinformatics analyses targeting the multinuclear nonheme iron oxidative (MNIO) enzyme family led to the identification of a *Streptomyces thermodiastaticus* JCM 4840 RiPP biosynthetic gene cluster, the *std* cluster, which includes multiple biosynthetic enzymes and a precursor peptide containing a conserved SNKEWQE motif. Using in vitro approaches, we elucidated the modifications installed by the *std* biosynthetic enzymes. First, a YcaO-TfuA pair thioamidates the backbone of asparagine. Next, a peptidase S8/S53 domain fused to a NodU-like carbamoyltransferase that both carbamoylates the ε-amino group of lysine to produce the non-proteinogenic amino acid homocitrulline and cleaves the C-terminal EWQE motif. Finally, a partner protein-MNIO pair bis-hydroxylates the β- and γ-carbon positions of the installed homocitrulline. The formation of homocitrulline and its subsequent modification are unprecedented in RiPP biosynthesis. Moreover, these findings expand the substrate scope of YcaO-TfuA enzymes and MNIOs and identify new roles for carbamoyl transferases in RiPP biosynthesis.

## INTRODUCTION

Ribosomally synthesized and post-translationally modified peptides (RiPPs) are a class of natural products originating from ribosomally produced precursor peptides that undergo post-translational modifications (PTMs) mediated by biosynthetic enzymes. These peptides are proteolytically processed and exported from the cell to fulfill their biological functions.^1-2^ The genetic framework encoding this process is organized into biosynthetic gene clusters (BGCs). The expansion of genomic sequencing initiatives across both cultured and uncultured organisms, coupled with advances in genome mining tools, has substantially contributed to the discovery of novel RiPP BGCs.^3-4^ Concurrently, the identification of new BGCs has emerged as a strategic approach for uncovering RiPP biosynthetic enzymes of previously unknown functions.^5^ Notably, RiPP BGCs enable direct functional assignment of biosynthetic enzyme activity, as the substrate, the precursor peptide, is typically encoded within the same BGC.^4^

The multinuclear nonheme iron oxidative enzyme family (MNIOs, formerly annotated as DUF692 proteins) is a class of RiPP biosynthetic enzymes that has attracted intense attention.^6-7^ To date, MNIOs have been shown to catalyze a range of unusual PTMs on proteinogenic amino acids within RiPP precursor peptides (Figure 1A) in the presence of iron and dioxygen, and, in some cases, require a partner protein to facilitate precursor peptide binding.^6-7^ PTMs of cysteine residues catalyzed by MNIOs were among the first to be characterized, including the formation of an oxazolone-thioamide in methanobactin,^8^ carbon excision in thiaglutamate,^9^ macrocycle formation in chryseobasin,^10^ and thiooxazole biosynthesis in the captophorins.^11-15^ Beyond cysteine, MNIOs can cleave the N–Cα bond in asparagine to generate a C-terminally amidated peptide,^16^ and modify aspartic acid, including conversion to a C-terminal α-keto acid and β-carbon hydroxylation.^17-19^ In addition, MNIOs catalyze bis-hydroxylation of phenylalanine residues during biophenomycin biosynthesis.^20^

**Figure 1.**
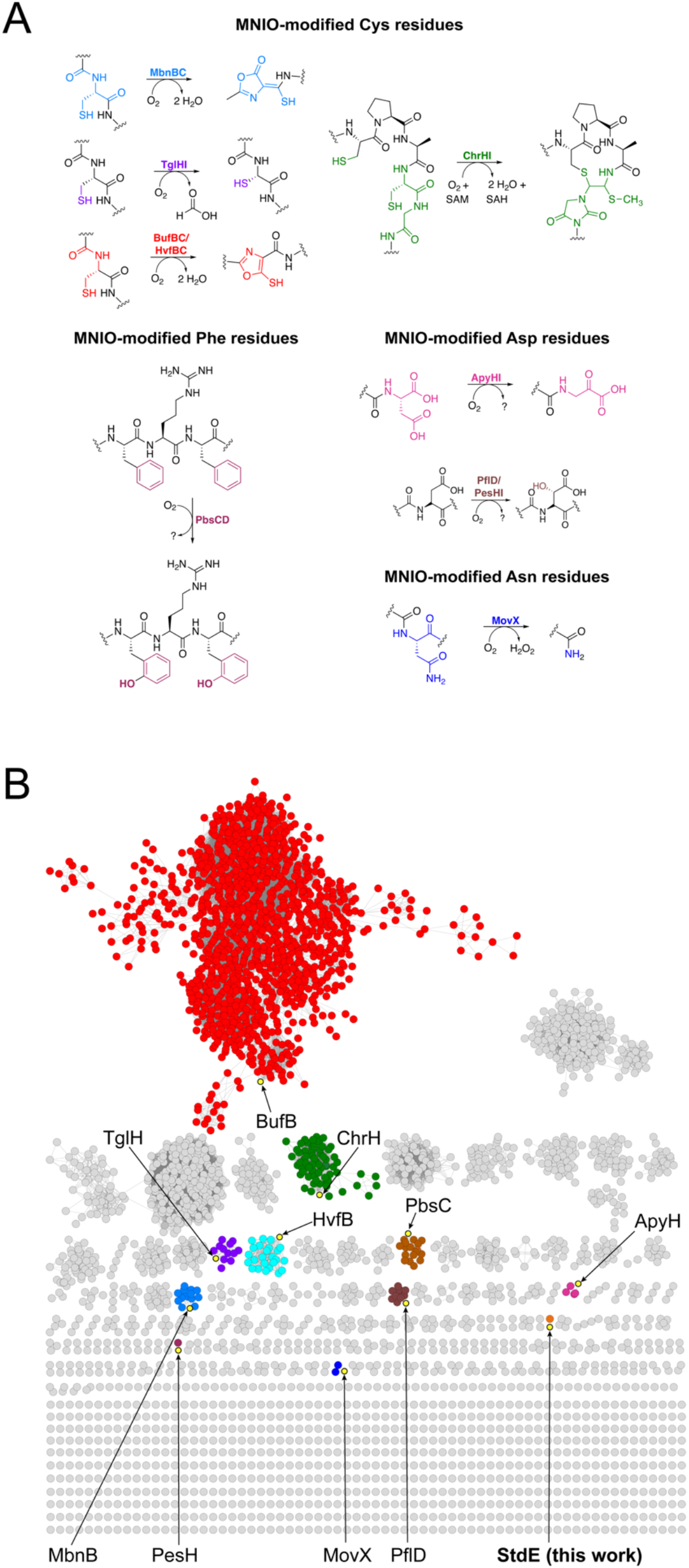
Functional diversity and bioinformatic analysis of the MNIO family. (A) Known PTMs catalyzed by MNIOs include modifications of cysteine, phenylalanine, asparagine, and aspartic acid residues. (B) Sequence similarity network (SSN) of the MNIO family comprising 13,864 members, generated using an SSN edge E-value of 5 and an alignment score threshold of 45. The RepNode 50 was visualized using the yFiles organic layout algorithm in Cytoscape v3.10. Experimentally characterized MNIOs are highlighted in yellow, with MNIO StdE emphasized as the focus of this study.

Motivated by the remarkable chemical diversity of reactions catalyzed by MNIOs, we sought to determine whether MNIOs can catalyze PTMs beyond those described previously. Using genome mining and a sequence similarity network (SSN), we identified an MNIO-containing BGC encoded in the genome of *Streptomyces (S*.*) thermodiastaticus* JCM 4840. This cluster, designated the *std* cluster, encodes multiple putative biosynthetic enzymes, including a partner–MNIO complex, a predicted peptidase S8/S53 domain fused to a NodU-like carbamoyltransferase, and a YcaO–TfuA pair. The corresponding precursor peptide contains a highly conserved C-terminal SNKEWQE motif. Here, using an in vitro approach, we show that these enzymes catalyze multiple unprecedented transformations of this motif. Specifically, the YcaO–TfuA pair catalyzes thioamidation of the Asn backbone, while the predicted peptidase S8/S53 domain fused to a NodU-like carbamoyltransferase generates the non-proteinogenic amino acid homocitrulline (Hcit) by carbamoylating the ε-amine of the Lys residue and cleaving the conserved EWQE residues. The partner–MNIO complex subsequently hydroxylates the β- and γ-carbons of the installed Hcit. These findings represent the first instance of a RiPP biosynthetic pathway that installs Hcit and expand the known catalytic scope of MNIOs and YcaO-TfuAs.

## RESULTS

### Identification of the *S. thermodiastaticus* JCM 4840 BGC

To identify MNIO-containing RiPP BGCs that encode unique conserved precursor peptide motifs, we generated a sequence similarity network (SSN) comprising approximately 13,864 MNIO sequences using the Enzyme Function Initiative (EFI) (Figure 1B).^21^ Notably, characterized MNIOs occupy only a small fraction of this network and are distributed across multiple clusters, highlighting the unexplored functional space within the MNIO family. MNIOs from uncharacterized clusters were prioritized, and their corresponding protein sequences were retrieved from UniProt and used to perform BLAST searches against the Integrated Microbial Genomes (IMG) database.^22^ This approach enabled direct inspection of MNIO-associated BGCs and their precursor peptides, thereby facilitating the identification of unusual precursor peptide sequences.

To identify an MNIO-containing BGC amenable to soluble heterologous expression in *E. coli* and to in vitro enzyme reactions, we focused on MNIO-containing BGCs from thermophilic bacteria, as other RiPPs have been successfully reconstituted using this approach.^23-28^ We selected an MNIO-containing BGC from the thermophilic strain *S. thermodiastaticus* JCM 4840, which we named the *std* cluster. The *std* cluster encodes a precursor peptide (StdA), a RiPP recognition element (RRE)-containing MNIO partner protein (StdD), an uncharacterized MNIO (StdE), a predicted peptidase S8/S53 domain fused to a NodU-like carbamoyltransferase (StdF), a YcaO enzyme (StdG), a TfuA protein (StdH), and an MFS-family permease (StdI) (Figure 2A, Table S1)). While MNIOs perform various PTMs (Figure 1A)^6-7^ and YcaO/TfuA pairs install backbone thioamides in precursor peptides,^29^ a peptidase S8/S53 domain fused to a NodU-like carbamoyltransferase has not been characterized in any context, including RiPP biosynthesis.^30^ To our knowledge, RiPPs from *S. thermodiastaticus* JCM 4840 have not been characterized previously, although the genus *Streptomyce*s is known to produce a wide variety of RiPPs.^31-33^

**Figure 2.**
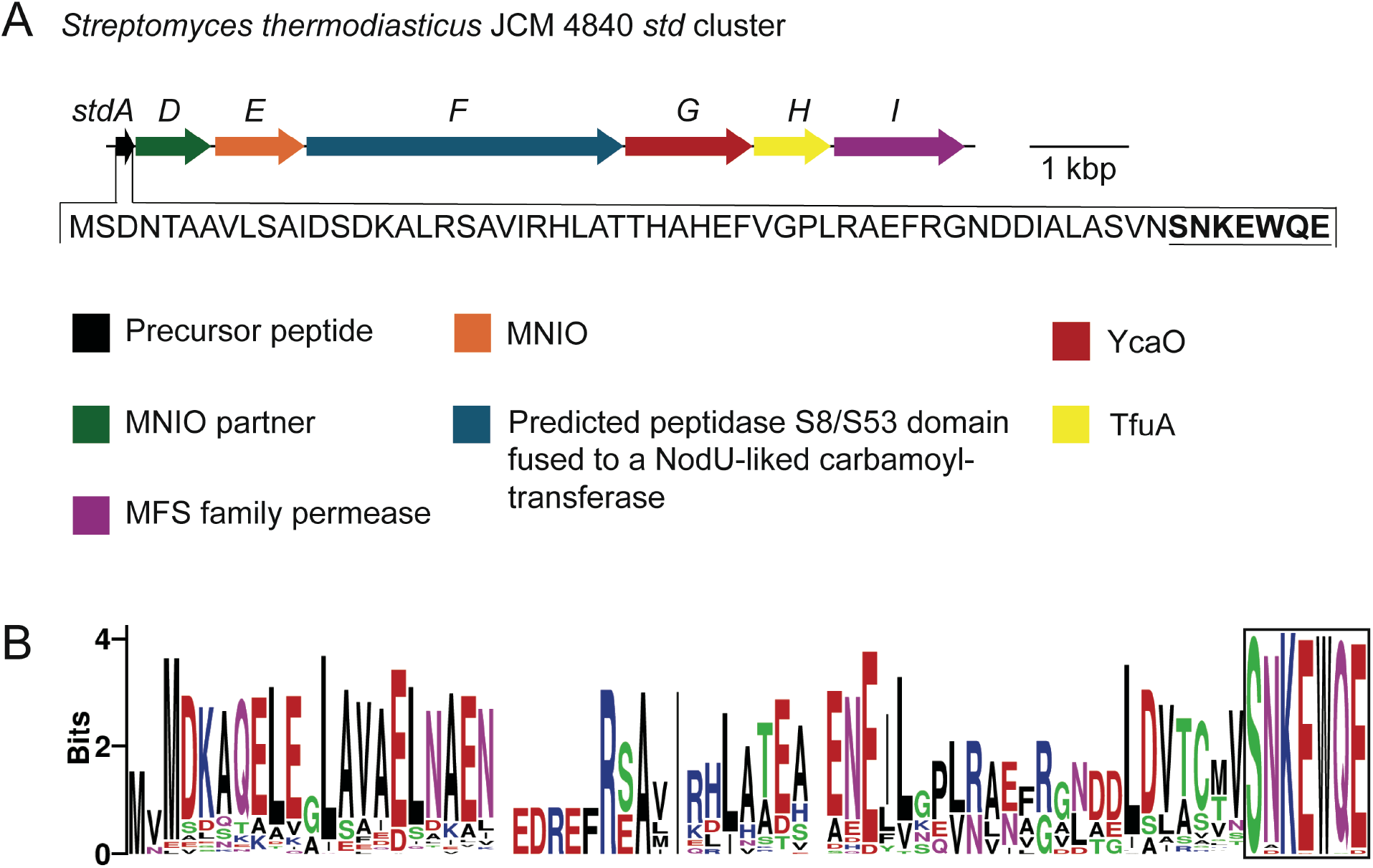
Analysis of the StdE-containing biosynthetic gene cluster from *S. thermodiastaticus* JCM 4840. (A) Organization of the *std* BGC with annotated genes. The predicted precursor peptide gene, *stdA*, is highlighted, and its amino acid sequence is shown. The predicted core peptide is bolded and underlined. (B) Sequence logo derived from the multiple sequence alignment of StdA and 96 orthologous precursor peptides, revealing a conserved C-terminal SNKEWQE motif (boxed).

To predict which StdA amino acids are modified by StdGH, StdF, and StdDE, we performed an IMG BLAST search using the StdA amino acid sequence, which only yielded 20 hits. We then performed an IMG BLAST search using the amino acid sequence of the MNIO, StdE, and manually identified 97 StdA ortholog hits (Table S2). The StdA orthologs were found across the Pseudomonadota, Actinomycetota, and Myxococcota phyla (Table S2 and Figure S1). Alignment of the precursor peptides revealed high conservation in the C-terminal region with 100% conservation of the C-terminal serine (S), lysine (K), and tryptophan (W) residues and 97-99% conservation of the two glutamic acid (E), asparagine (N), and glutamine (Q) residues (Figure 2B). As the most highly conserved region of StdA, we hypothesized that the SNKEWQE motif is the core peptide likely to be modified by StdGH, StdF, and/or StdDE. This predicted core peptide does not resemble any other known core peptides modified by MNIOs.

### Assignment of Std Enzyme Activities and Order of Modifications

With three enzymes encoded in the *std* cluster, StdGH, StdF, and StdDE, the identity of the first biosynthetic step was not obvious. Since previously characterized MNIOs can modify unmodified precursor peptides,^8,10-14,16,20^ we initially hypothesized that StdDE catalyzes the first modification of StdA. To test whether StdDE directly modifies StdA, *stdD* and *stdE* were codon-optimized for heterologous expression in *E. coli* and co-expressed, as complex formation of MNIOs with their partner proteins typically enhances solubility. Using an N-terminal hexahistidine tag on StdD, the StdDE complex was purified by cobalt-affinity chromatography followed by size-exclusion chromatography, eluting as a single peak consistent with stable complex formation (Figure S2A). Iron quantification by Inductively Coupled Plasma Mass Spectroscopy (ICP-MS) revealed 2.2 iron atoms per StdDE heterodimer, reflecting partial incorporation of the metal cofactor.^34-35^ To maximize iron loading and potential catalytic activity, StdDE was incubated overnight at room temperature with unmodified StdA in the presence of freshly prepared ammonium iron(II) sulfate hexahydrate and ascorbate.^34-35^ The reaction was then heated to precipitate StdDE and to isolate the potentially modified peptide for analysis. However, no mass change was detected by Liquid Chromatography Time-of-Flight Mass Spectrometry (LC-TOF-MS) (Figure 3i, ii), suggesting that premodification of StdA might be required for reaction with StdDE.

**Figure 3.**
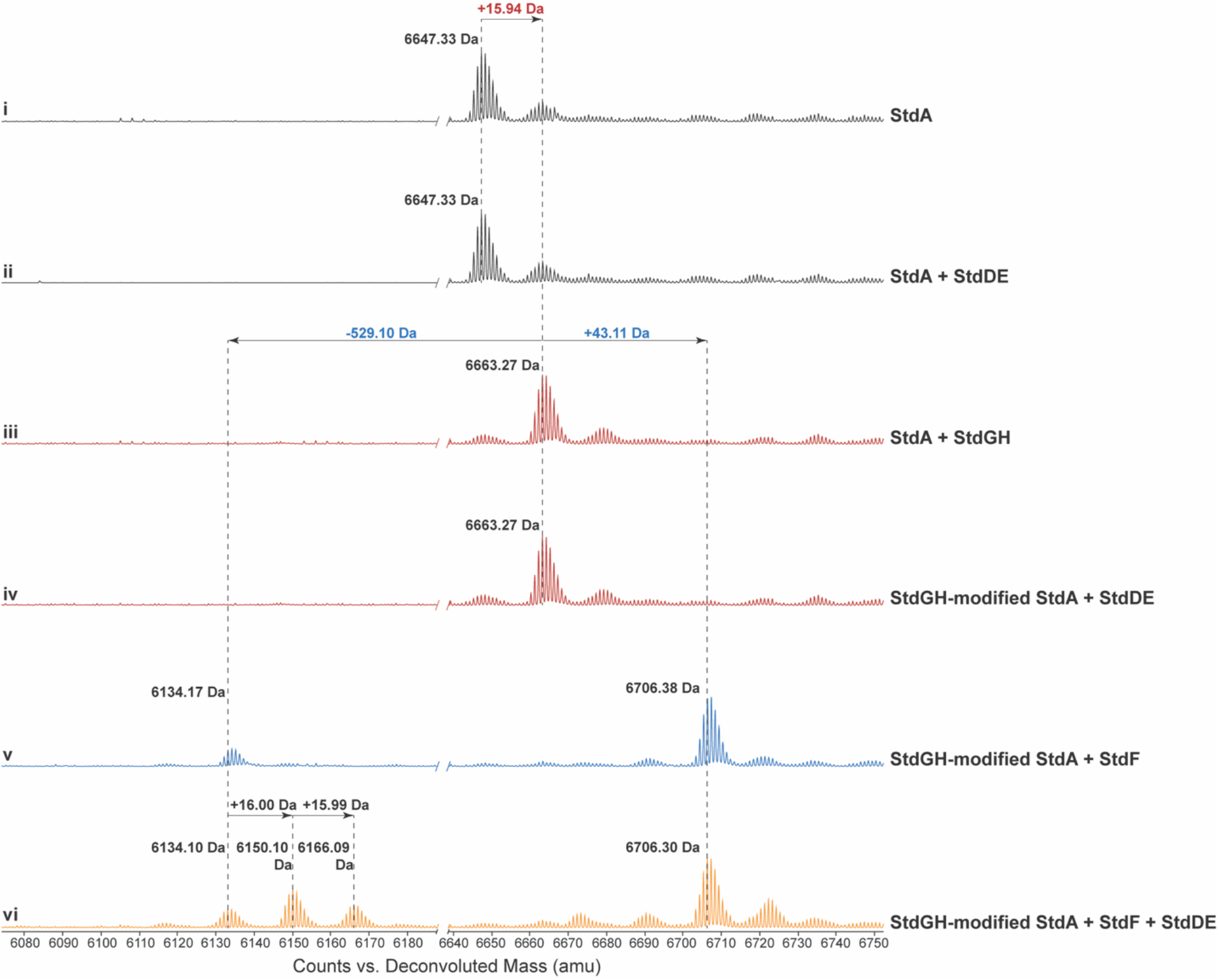
LC-TOF-MS analysis of in vitro reconstitution of StdGH, StdF, and StdDE to determine the substrate specificity and activity of StdDE. ESI-MS deconvoluted spectra of (i) the StdA precursor peptide alone, (ii) after incubation with StdDE, and (iii) after incubation with StdGH. The StdGH-modified StdA peptide was subsequently incubated with (iv) StdDE and (v) StdF. The StdGH-modified StdA peptide was then incubated with both (vi) StdF and StdDE.

The presence of StdGH, a predicted YcaO–TfuA pair, prompted the hypothesis that thioamidation of StdA constitutes the initial biosynthetic step. YcaO–TfuA pairs catalyze backbone thioamidation in an ATP- and sulfur-dependent manner and often act as gatekeepers in RiPP biosynthesis.^36-39^ To test this hypothesis, *stdG* and *stdH* were codon-optimized and co-expressed in *E. coli* with an N-terminal hexahistidine tag on StdH. The StdGH complex was purified by cobalt-affinity and size-exclusion chromatography, yielding a single peak (Figure S2B). Overnight incubation of StdA with StdGH in the presence of ATP and Na_2_S. The reaction was then heated to remove StdGH and the resulting peptide harbored a 15.94 Da mass gain detected by LC–TOF–MS, consistent with backbone thioamidation (Figure 3iii).^37^ We then incubated the StdGH-modified StdA with StdDE overnight in the presence of freshly prepared ammonium iron(II) sulfate hexahydrate and ascorbate, and again observed no detectable mass shift of the peptide (Figure 3iv).

We next investigated the function of the predicted peptidase S8/S53 domain fused to a NodU-like carbamoyltransferase, StdF. StdF was expressed and purified analogously to StdDE and StdGH, yielding soluble protein (Figure S2C, D). Following protocols used for the carbamoyl transferase, GdmN,^40^ StdGH-modified StdA was incubated overnight with StdF in the presence of carbamoyl phosphate and ATP followed by heating to remove StdF. LC-TOF-MS of the peptide revealed two mass changes: a 43.11 Da mass gain that is consistent with carbamoyl group installation, and a 529.10 Da mass loss predicted to correspond to carbamoylation and proteolytic removal of the C-terminal EWQE sequence, likely mediated by StdF’s predicted peptidase S8/S53 domain (Figure 3v).

With StdDE remaining as the final biosynthetic enzyme with no assigned function in the *std* cluster, the StdGHF-modified peptides were incubated with StdDE in the presence of freshly prepared ammonium iron(II) sulfate hexahydrate and ascorbate overnight and the reaction was heated to precipitate and remove StdDE. Surprisingly, no mass gain or loss of the peptide was detected by LC-TOF-MS (Figure S3). We reasoned that the additional heating step used to inactivate and remove StdF may have compromised StdGHF-modified StdA integrity. To directly supply the StdGHF-modified peptides for StdDE, StdGH-modified StdA was incubated overnight with StdF in the presence of ATP, carbamoyl phosphate, StdDE, ammonium iron(II) sulfate hexahydrate, and ascorbate. This reaction produced a peptide with two mass gains of 16.00 Da and 15.99 Da on the StdF cleaved product (Figure 3vi), consistent with two hydroxylation events.

### Localization of the Modifications

To localize the mass gains and losses resulting from the in vitro enzymatic reactions, we digested the products of unmodified StdA, StdGH-modified StdA, StdGHF-modified StdA, and StdGHFDE-modified StdA with Arg-C Ultra endopeptidase. Arg-C Ultra was chosen because it generates fragments containing the predicted core motif, SNKEWQE (Figure 4A). The resulting fragments were purified by HPLC (Table S11) and analyzed by LC-TOF-MS to confirm their masses (Figure S4). Arg-C Ultra digestion yielded 18-mer peptides from unmodified StdA, StdGH-modified StdA, and StdGHF-modified StdA, whereas 14-mer peptides were obtained from StdGHF- and StdGHFDE-modified StdA. The shorter peptides are consistent with the predicted proteolytic cleavage of the EWQE motif by StdF.

**Figure 4.**
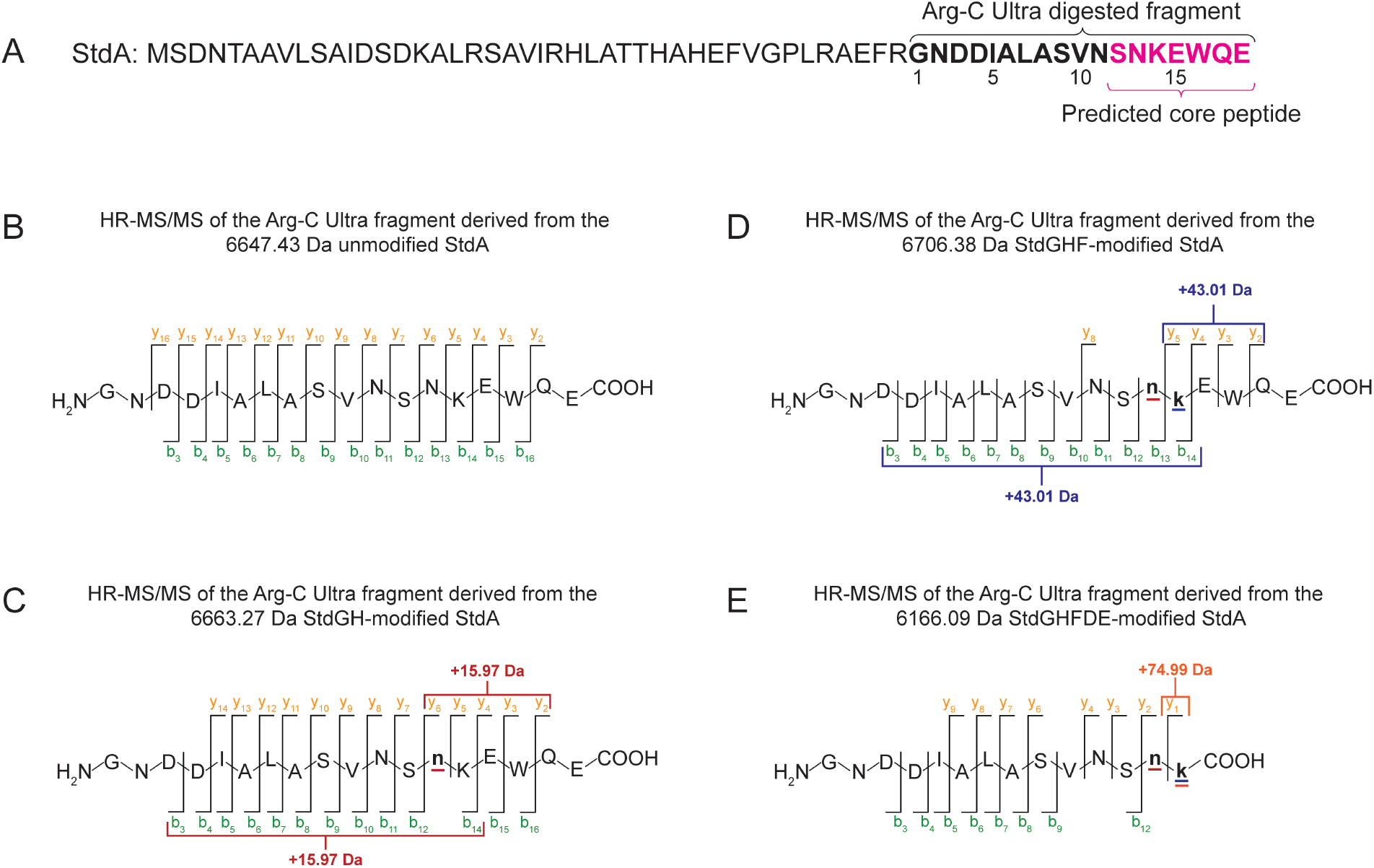
Localization of modifications. (A) Amino acid sequence of StdA; the Arg-C Ultra cleavage site generating the predicted core fragment is highlighted in magenta. (B-E) HR-MS/MS fragmentation patterns of the corresponding digested peptides: (B) unmodified, (C) StdGH-modified, (D) full-length StdGHF-modified, and (E) StdGHFDE-modified. b- and y-ion values are listed in Tables S3-6.

High-resolution tandem mass spectrometry (HR-MS/MS) was used to identify the modification sites within the Arg-C Ultra digested fragments (Figures S5-S8). The Arg-C Ultra digested fragment of unmodified StdA served as a control (Figure 4B). Analysis of Arg-C Ultra digested StdGH-modified StdA revealed the 15.97 Da mass gain corresponding to a predicted thioamide installation, localized to Asn13 within the conserved SNKEWQE motif (Figure 4C). HR-MS/MS analysis of Arg-C Ultra digested StdGHF-modified StdA localized the 43.01 Da mass gain to Lys14 (Figure 4D). This mass gain led to our hypothesis that a carbamoyl group is installed on the Lys14 ε-amino group, producing the non-proteinogenic amino acid, Hcit. Surprisingly, HR-MS/MS analysis of Arg-C Ultra digested StdGHFDE-modified StdA revealed a 74.99 Da mass gain localized to Lys14 (Figure 4E) and the absence of the EWQE motif, suggesting again the production of Hcit by StdF, the cleavage of the EWQE motif by StdF, and the installation of two hydroxyl groups on the proposed Hcit by StdDE.

### Structural elucidation of StdGHFDE-modified ^15^N/^13^C-Asn13-Lys14-StdA

Having established that only the Asn13 and Lys14 residues within the SNKEWQE motif are modified, we predicted that Asn13 harbors a backbone thioamide, the Lys14 is converted to an Hcit and harbors two hydroxyl groups, and the EWQE segment is proteolytically removed. To determine the structure of the StdGHFDE-modified StdA product, we conducted nuclear magnetic resonance (NMR) analysis of the PTMs. We obtained isotopically labeled StdA in which the Asn56 and Lys57 residues were universally enriched with both ^13^C and ^15^N. Large-scale enzymatic reactions with the labeled StdA yielded a final StdGHFDE-modified product 13.99 Da heavier than the reactions with the unmodified peptide (Figure S9), indicating full incorporation of ^13^C and ^15^N at Asn56 and Lys57. Arg-C Ultra digestion, as described above, was used to generate an unmodified ^15^N/^13^C-Asn13-Lys14-StdA and a StdGHFDE-modified ^15^N/^13^C-Asn13-Lys14-StdA, in which Asn13 and Lys14 correspond to Asn56 and Lys57 in the intact StdA (Figures S10-11).

A classic suite of 2D NMR assignment experiments was collected at natural abundance for both the unmodified ^15^N/^13^C-Asn13-Lys14-StdA and the StdGHFDE-modified ^15^N/^13^C-Asn13-Lys14-StdA, consisting of [^1^H, ^1^H] Total Observed Correlation SpectroscopY (TOCSY) and Rotational Overhauser Exchange SpectroscopY (ROESY) homonuclear experiments and respective ^15^N or ^13^C Heteronuclear Single Quantum Correlation (HSQC) experiments (Figures S12-13). Connectivity patterns of the TOCSY were used to identify potential residue systems, and the heteronuclear connectivity of the HSQCs were used to assign specific atom types to the TOCSY patterns. Sequential connectivity of the isolated spin systems was then achieved via analysis of ROESY spectra (Figure S12C). The spin systems for Asn13 and Lys14 were then identified, with a large downfield movement of the Lys14 amide chemical shift.

The Correlation Observed SpectroscopY (COSY) spectral pattern of the resonances associated with the anomalously deshielded amide proton is shown in Figure 5A. This pattern putatively matches that of lysine, but shows two key anomalies, namely the shifts of the H^b^ and H^g^ nuclei. In a typical lysine, these nuclei resonate between 2 and 1 ppm like other aliphatic protons removed from electron withdrawing groups. In this spin system, both are found between 4 and 3 ppm, significantly downfield of where they would normally be, but consistent with the introduction of a geminal hydroxyl group at each position. The COSY pattern confirms the identify of these modified resonances through limited spin (3-bond) coupling as compared to the TOCSY. The H^a^ signal only sees the H^b^, the H^b^ sees the H^a^ and H^g^ only, and so on. Analysis of the enriched ^13^C-HSQC in Figure 5B shows that the carbon atoms directly attached to the H^b^ and H^g^ nuclei are shifted 40 ppm downfield relative to their unmodified counterparts as well, also consistent with electron withdrawing hydroxyl attachment.

**Figure 5.**
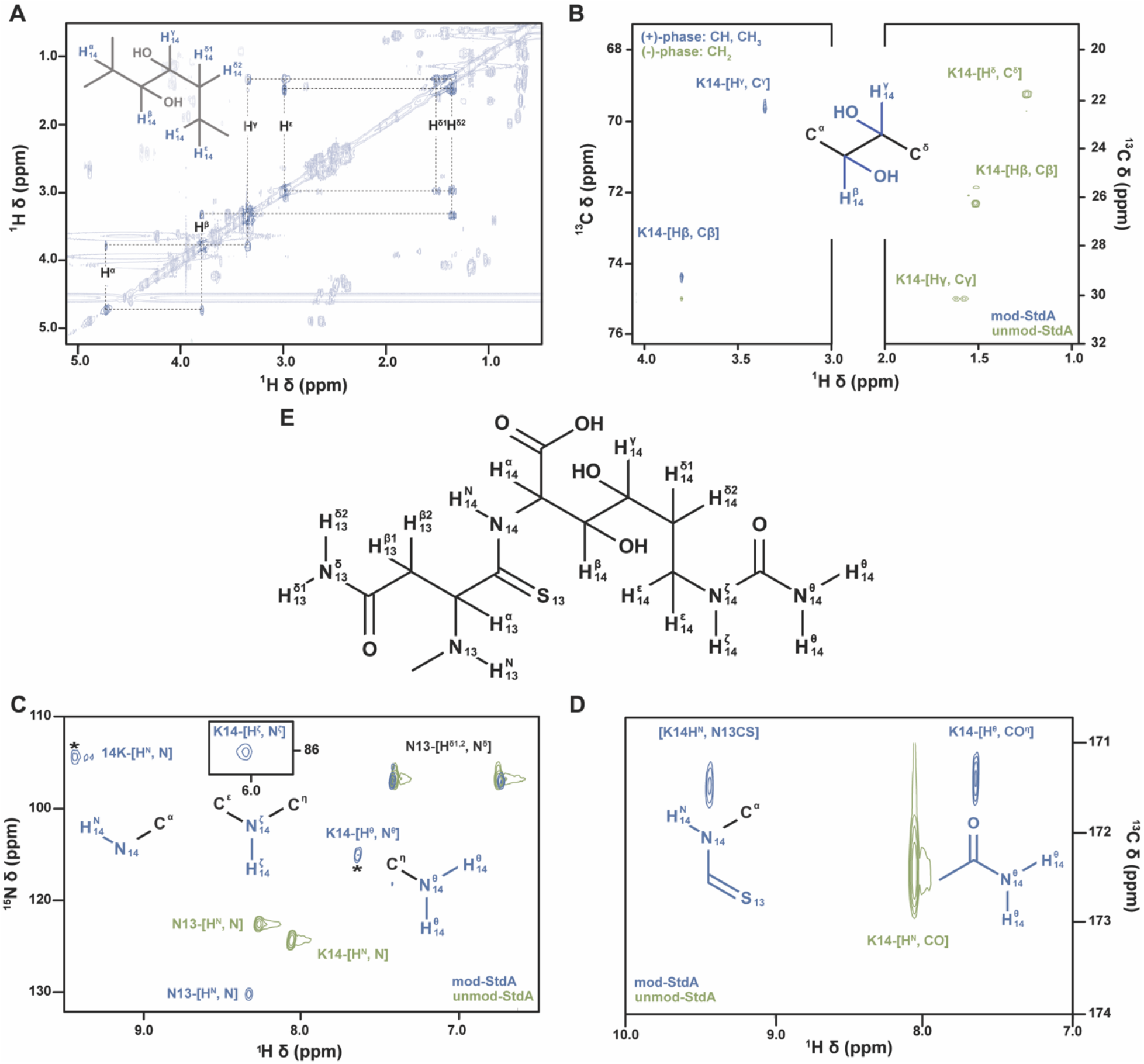
Structure determination of Arg-C Ultra digested StdGHFDE-modified ^15^N/^13^C-Asn13-Lys14-StdA. All modified spectra in this figure are in blue while unmodified spectra are in green. (A) Aliphatic region of the StdGHFDE-modified ^15^N/^13^C-Asn13-Lys14-StdA COSY spectrum, highlighting sequential resonance contacts within the Lys14 spin system in bolded contours. (B) Composite of modified and unmodified ^13^C-HSQCs. Phase of protons geminal to position of hydroxylation show clear phase change upon modification (green to blue). (C) ^15^N-HSQC of modified and unmodified peptides showing the formation of two new peaks and the ∼1 ppm shift of the Lys14 H^N^. Peaks with asterisks are at lower contour levels due to contaminating T_1_ noise from differential isotopic enrichment. (D) Pseudo-2D HNCO spectra of modified and unmodified StdA showing the formation of a new N-CO correlation and the movement of the Lys14 H^N^. (E) Proposed StdGHFDE-modified StdA structure.

The enriched ^15^N-HSQCs in Figure 5C reveal not only the expected Asn13 backbone, Asn13 sidechain, and the Lys14 backbone [^1^H-^15^N] amide pairs, but the presence of two additional peaks at (7.6, 117.6) and (6.0, 87.2) ppm in the modified ^15^N/^13^C-Asn13-Lys14-StdA spectrum. To additionally characterize these new amide protons, we collected HNCO experiments of the isotopically enriched, modified and unmodified StdA peptides and observed the formation of a new peak at (7.6, 171) in the modified StdA HNCO (Figure 5D). The carbonyl shift of this new peak and the Lys14 backbone are close, yet substantially different, suggesting these two amide groups are connected to different carbonyl-like groups. The proton at 6.0 ppm likely does not show a signal in the HNCO due to a difference in ^1^J coupling preventing effective magnetization transfer.

The backbone amide proton of the terminal Lys14 shows a greater than 1 ppm difference in chemical shift between the modified and unmodified StdA (Figure 5C). This major perturbation of the normal amide electronic environment suggests a deshielding effect from neighboring chemical groups. Additionally, the C^α^ and H^α^ of the modified lys14 show significant chemical shift perturbations consistent with the desheidling effects of a more electron withdrawing group attached to the amide (Figure S13). Formation of a thioamide backbone linkage between Asn13 and Lys14 is consistent with these observed spectral features due to the increased radius and electron density of the sulfur further depolarizing the carbonyl carbon.

Taken together, these NMR data confirm the StdE-catalyzed hydroxylation of the β- and γ-carbons of the terminal Hcit, the StdF-catalyzed carbamoylation of the terminal Lys ζ-nitrogen to create Hcit, and the StdGH-catalyzed thioamidation of the proceeding Asn13 carbonyl to form a thioamide backbone linkage between Asn13 and Lys14. The proposed structure is shown in Figure 5E. Protons geminal to the site of hydroxylation in typical amino acids (Ser, Thr) show a significant downfield shift, as do the shifts of the attached carbons, both of which are observed for the β- and γ-carbons herein. The appearance of the ζ proton resonance is consistent with modification slowing the amide-solvent exchange rate, which renders this signal unobservable except at very acidic pHs for typical lysines. Combined with the appearance of a new peak in the HNCO, the most likely modifying group is a carbamoyl. The movement of the Lys14 backbone amide ∼1 ppm downfield is consistent with the presence of a more electron-withdrawing, larger atom, like sulfur, replacing the Asn13 carbonyl oxygen. The larger electron cloud creates an anisotropic deshielding effect, moving the shift downfield.

## DISCUSSION

Here we have elucidated the RiPP biosynthetic pathway encoded by the *std* BGC from *S. thermodiasticus* JCM 4840 (Figure 6). Biosynthesis is initiated by StdGH, which replaces the backbone carbonyl oxygen of an Asn residue with sulfur, forming a thioamide. StdG likely follows the canonical YcaO mechanism, catalyzing thioamide formation through ATP-dependent activation of the peptide backbone, while StdH likely functions analogously to TfuA, delivering sulfur to the activated intermediate to complete thioamide installation.^36,41^ The incorporation of a thioamide at Asn may confer enhanced protease resistance or backbone rigidity on the final product, consistent with the known role of thioamides in stabilizing peptide conformation in other RiPP natural products.^42^ YcaO-TfuA-modified RiPPs characterized to date fall into four classes: thioviridamide-like,^38,43-44^ thiovarsolins,^45^ and thioamidated thiopeptides,^46-47^ with chimeric lanthipeptides recently added as an additional class.^48^ None of these characterized RiPPs contain a YcaO-TfuA-modified Asn residue, and the StdGHFDE-modified StdA does not fall into any of the existing classes. Therefore, the StdGH reaction expands the substrate scope of YcaO-TfuA enzymes and defines a new class of YcaO-TfuA-modified RiPPs.

**Figure 6.**
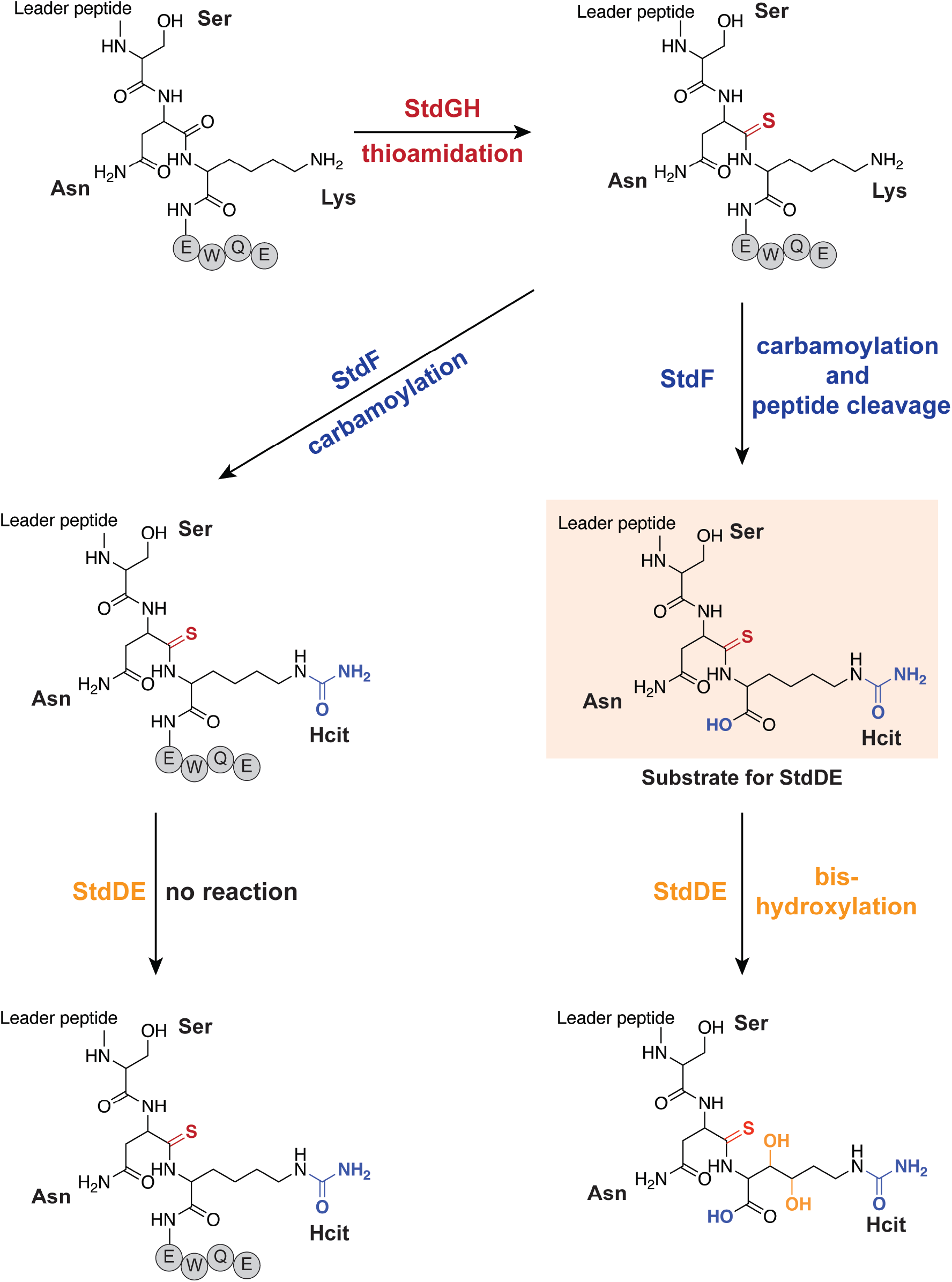
Summary of enzymatic activities of the *std* cluster enzymes and proposed structures. StdGH catalyzes the thioamidation of the Asn backbone carbonyl. StdF then carbamylates the ε-amino group of the Lys to produce Hcit and performs proteolytic cleavage of the EWQE motif, which is required for selective hydroxylation of β- and γ-carbon positions of Hcit by StdDE.

In the next biosynthetic step, StdF acts bifunctionally: it carbamoylates the conserved Lys ε-amine to produce Hcit, a rare non-proteinogenic amino acid not previously observed in natural product pathways, and removes the terminal EWQE sequence by proteolysis, thereby generating the substrate for StdDE. The carbamoyl transferase domain of StdF likely operates via a mechanism analogous to that of characterized carbamoyl transferases, involving an ATP-dependent transfer of a carbamoyl group from carbamoyl phosphate to an amino group.^40^ While carbamoyl transferases have been implicated in the biosynthesis of many bioactive secondary metabolites, their involvement in RiPP biosynthesis is exceedingly rare.^49-52^ Indeed, crocagin is the only well-characterized RiPP whose biosynthesis involves a carbamoyl transferase.^53-54^ Thus, StdF represents the second example of a carbamoyl transferase associated with RiPP biosynthesis and the first RiPP-associated enzyme to catalyze the carbamoylation of lysine ε-amine to produce Hcit.

Hcit is primarily known to arise from the nonenzymatic reaction of isocyanic acid with lysine residues or protein N-termini. The only indirect evidence for enzymatic formation is a mitochondrial ornithine carbamoyltransferase-like activity proposed to convert lysine and carbamoyl phosphate to Hcit.^55-58^ Importantly, Hcit should not be confused with citrulline, which differs by a single methylene unit. Citrulline-containing RiPPs are generated from arginine deimination catalyzed by bacterial peptidylarginine deiminases,^59^ but no such enzymes are encoded within the *std* cluster. These observations further support StdF-catalyzed carbamoylation of lysine as the source of Hcit, and establish StdF as the first enzyme to biosynthesize this PTM in a in a natural product pathway, in contrast to its exclusively non-enzymatic, pathology-associated origins in human biology.^55-56^

StdF also catalyzes the proteolytic removal of the C-terminal EWQE sequence, likely following carbamoylation, thereby generating the substrate for StdDE. While the SNKEWQE motif was initially predicted as the full core peptide based on sequence conservation, the proteolytic activity of StdF reveals that the functional core is limited to the SNK tripeptide, with EWQE serving as a C-terminal follower peptide. This activity is presumably mediated by the predicted peptidase S8/S53 domain of StdF. Removal of follower peptides is common in RiPP biosynthesis. For example, PatG cleaves C-terminal follower peptides during cyanobactin maturation, PCY1 processes the C-terminal sequence to generate segetalin A, and SpyE excises C-terminal peptides to produce peptide–nucleobase hybrids.^60-62^ The fusion of a follower peptide peptidase S8/S53 domain to a carbamoyl transferase in StdF is unique. Further mechanistic studies will be required to investigate the active sites of both domains and to establish the precise order of these modifications.

Finally, StdDE catalyzes bis-hydroxylation at the β- and γ-carbon positions of the newly installed Hcit residue of the StdGHF-modified peptide without the EWQE motif. MNIOs have been reported to bis-hydroxylate Phe residues and singly hydroxylate Asp residues,^18,20^ but hydroxylation of non-proteinogenic amino acids has not been observed previously, expanding the substrate scope of MNIOs. We propose a mechanism analogous to β-carbon hydroxylation of Asp by the MNIO PflD, for which an Fe(IV)-oxo intermediate has been suggested, though not yet experimentally established, to perform C–H activation.^18^ An AlphaFold 3 model of StdDE bound to StdA positions the Lys side chain in proximity to the predicted trinuclear iron active site (Figure S14), providing a structural rationale for the observed selectivity. Removal of the EWQE motif probably allows for positioning of the substrate. Further mechanistic and spectroscopic studies will be needed to characterize the iron cluster of StdDE and establish the stereochemical outcome of each hydroxylation of Hcit.

## CONCLUSION

Through in vitro reconstitution of the *std* cluster biosynthetic enzymes, we have established the sequential modification of the precursor peptide SNKEWQE core motif: backbone thioamidation of Asn by a YcaO–TfuA pair (StdGH), enzymatic generation of Hcit and peptide cleavage by a peptidase S8/S53 domain fused to a NodU-like carbamoyltransferase (StdF), and bis-hydroxylation of the non-proteinogenic amino acid Hcit by an MNIO (Figure 6). Among these modifications, the enzymatic installation of Hcit is perhaps the most unexpected. In humans, Hcit arises exclusively through non-enzymatic carbamoylation of lysine, a pathological modification linked to aging, inflammation, and rheumatoid arthritis,^55-56^ yet the *std* cluster uses a dedicated enzyme, StdF, to install the same modification site-specifically and to use it as a substrate for an MNIO. Thus, Hcit, long viewed as a marker of chemical damage in human biology, can serve as a RiPP building block. The identity of the *std* cluster mature RiPP and the protease responsible for leader peptide removal remain under investigation, as does the biological activity of the mature RiPP. Together, these results underscore the power of MNIO genome mining to discover RiPP BGCs that produce unprecedented chemical modifications

## MATERIALS AND METHODS

### Materials

All chemicals were used without further purification unless stated otherwise. All antibiotics, media, and inducers were purchased from Research Products International (RPI). Codon-optimized plasmids for StdA, StdDE, StdF, and StdGH expression were purchased from GenScript Biotech. Arg-C Ultra protease was purchased from New England Biolabs (NEB). Homocitrulline was purchased from Millipore Sigma. Peptide synthesis and purifications were performed by the Peptide Synthesis Core Facility of the Center of Regenerative Nanomedicine at Northwestern University. DNA and amino acid sequences are included in the Supporting Information.

### General procedures

Heterologous expression of enzymes was performed using an Innova® 44 incubator shaker. Protein purification was performed using TALON Metal Affinity Resin and an AKTA pure™ chromatography system with a Superdex 200 Increase SEC column, 10/300 GL (Cytiva). Protein and peptide concentrations were obtained using a NanoDrop One spectrophotometer. High-performance liquid chromatography (HPLC) was performed on an Agilent 1100 series HPLC system equipped with a diode-array absorbance detector from 190 to 950 nm, Peltier-thermostatted auto-sampler (4-40 °C), programmable automatic injector for 20-900μl sample, vacuum degasser, Peltier-thermostatted column compartment (4-80 °C), and a quaternary pump system using a high-speed proportioning valve, located in the Northwestern Keck Biophysics Facility. High-resolution mass spectrometry (HR-MS/MS) was performed by the Northwestern Proteomics Core Facility using an Orbitrap Exploris 240. Data were collected for 60 min in data-dependent acquisition mode. Data analysis was performed using FragPipe. Nuclear magnetic resonance (NMR) spectra were collected at the Integrated Molecular Structure Education and Research Center (IMSERC) at Northwestern. NMR spectra were collected using a Bruker Neo 600 MHz system with QCI-F cryoprobe which was operated using the Bruker TopSpin software. 1D/2D NMR data were analyzed with MestReNova, and figures were generated in Adobe Illustrator.

### Sequence similarity network analysis

The sequence similarity network (SSN) of MNIOs was created using the Enzyme Function Initiative Enzyme Similarity Tool (EFI-EST) web resource (https://efi.igb.illinois.edu/efi-est/). The MNIO Pfam (Pfam ID: PF05114) was used as a “family” query to generate SSNs for all MNIO proteins identified in UniProtKB. As of June 2025, the MNIO family comprised 13,864 members. Additionally, an E-value of 5 was used for the SSN edge calculation. An alignment score of 45 was applied to create the final SSN. The representative node 50 was visualized using the yFiles organic layout algorithm, Cytoscape v3.10 add-in.

### Alignment of StdA orthologs

StdA and its precursor peptide orthologs were aligned using MultAlin (http://multalin.toulouse.inra.fr/multalin/). The sequence logo was created using WebLogo.(https://weblogo.berkeley.edu).

### Unmodified StdA preparation

The lyophilized peptide was dissolved in a buffer containing 50 mM HEPES pH 8, 150 mM KCl, and 10% glycerol to a final concentration of 1 mM or 2 mM, flash-frozen, and stored at -80 °C for later use.

### Heterologous expression of StdDE, StdGH, and StdF

The StdE, StdF, StdG, and StdH genes were codon-optimized for *E. coli* using GenScript’s optimization tool, then synthesized and cloned by GenScript into a pMAL-c5x vector or pRSFDuet vector (Table S7). The pRSFDuet vector was chosen for its ability to co-express two genes, which is beneficial because StdD and StdE, as well as StdG and StdH, are predicted to form complexes.

For protein expression, plasmids were transformed into *E*. coli BL21 electrocompetent cells via electroporation for pRSFDuet_StdDE, pRSFDuet_StdF, or into Rosetta2 (DE3) pLysS chemically competent cells for pRSFDuet_StdGH. For pRSFDuet_StdDE and pRSFDuet_StdF, transformation, 2 μL of DNA was added to 50 μL of electrocompetent *E. coli* BL21 cells. The cells were then pipetted into a GenePulser® Cuvette (0.1 cm electrode gap) and electroporated using a Bio-Rad MicroPulser^™^. The transformed cells were recovered in 1 mL of warmed S.O.C. medium (ThermoFisher, Invitrogen) and shaken at 37 °C for 1 h at 200 rpm. The recovered cells were then plated onto LB-agar plates containing the appropriate antibiotic (Table S7) The plates were incubated at 37 °C overnight.

For the pRSFDuet_StdGH transformation, 2 μL of DNA was added to 50 μL of chemically competent *E. coli* Rosetta2 (DE3) pLysS cells. The cells were heat shocked at 42 °C for 4 s on a heating block. Afterward, they were placed on ice for 2 min, then recovered in 1 mL of S.O.C. medium with shaking at 37 °C for 1 h at 200 rpm. The recovered cells were then plated onto LB-agar plates containing the appropriate antibiotics (Table S7). These plates were incubated overnight at 37 °C.

To express StdDE, StdGH, or StdF, a single transformed colony was added to 50 mL of sterile LB media containing the appropriate antibiotics for the plasmid and strain. The culture was incubated overnight at 37 °C with shaking at 200 rpm. The next day, 10 mL of this overnight culture was transferred into 1 L of sterile LB (for StdGH) or 1 L of sterile TB (for StdF, and StdDE). The flask was shaken at 37 °C at 200 rpm until the optical density at 600 nm (OD_600_) reached 0.6. Afterwards, the cultures were cooled on ice for 30 min. For StdDE, ammonium iron(II) sulfate hexahydrate was added to reach a final concentration of 300 μM. Once cooled, IPTG was added to the culture at a final concentration of 200 μM to induce protein expression. The culture was then shaken overnight at 18 °C and 180 rpm. The next day, the cell pellet was obtained by centrifuging the culture in a 1 L plastic tube at 10000 × g at 4 °C for 10 minutes. The supernatant was discarded, and the pellet was collected and stored at -80 °C for future purification.

### Purification of StdDE, StdGH, and StdF

The frozen cell pellet was thawed in Buffer A (50 mM HEPES pH 8, 400 mM KCl, 10% glycerol) with one tablet of cOmplete^™^, EDTA-free Protease Inhibitor Cocktail (Millipore Sigma). The remaining steps were performed at 4 °C. After resuspending the cells in Buffer A, the mixture was transferred to a metal beaker, and an ultrasonic processor sonicator horn was placed into the mixture to lyse the cells at 30% amplitude, with 1 s on and 3 s off for 4 min. The total cell lysate was then centrifuged at 70000 × g for 1 hour at 4 °C using the ultracentrifuge. The supernatant was then applied to Buffer A-equilibrated TALON cobalt resin (Takara Bio). After the supernatant was loaded, the resin was washed with five column volumes of Buffer A, and the bound enzyme was eluted with five column volumes of Buffer B (50 mM HEPES pH 8, 400 mM KCl, 250 mM imidazole, 10% glycerol). The collected protein was then concentrated to 2 mL using a Millipore Amicon Ultra centrifugal filter, 30 kDa molecular weight cut-off and a tabletop centrifuge at 3000 × g. The 2 mL concentrated protein solution was then applied to a Buffer C (50 mM HEPES pH 8, 150 mM KCl, 10% glycerol)-equilibrated S200 column for further purification by size-exclusion chromatography on an AKTA system. The protein purity was assessed by SDS-PAGE (Figure S2D), and the protein was flash-frozen in liquid nitrogen and stored at -80 °C.

### ICP-MS sample preparation

ICP-MS samples were prepared in 5% trace metal-free nitric acid and 1% trace metal-free hydrogen peroxide, with 5 ppb of indium, lithium, scandium, and yttrium standards (Inorganic Ventures). The concentrations of iron, cobalt, nickel, copper, and zinc were determined using the NWU-16 multielement standard (Inorganic Ventures).

### Intact Mass Spectrometry

LC-TOF-MS analyses were performed using an Agilent 6230 LC-TOF mass spectrometer operated in positive-ion ESI mode and coupled to an Agilent 1200 Series HPLC system equipped with a binary pump and an Agilent 1200 autosampler. Peptides were separated on a 10 cm C18 column maintained at 35 °C. The mobile phase consisted of water with 0.1% formic acid (buffer A) and acetonitrile with 0.1% formic acid (buffer B). The chromatographic method was as follows: 100% A (0-0.5 min, 0.4 mL/min), 0-100% B (0.5-5.0 min), 100% B (5.0-7.25 min), return to 100% A (7.25-7.5 min, flow increased to 0.5 mL/min), followed by equilibration at 100% A (7.5-10.0 min). Data were analyzed using Agilent MassHunter BioConfirm 10.0. Chromatographic peaks were selected at their maximum height to extract mass spectra. Mass deconvolution was performed using the maximum entropy algorithm with an m/z range of 500-2,000 (Tables S8-S10), an expected mass range of 6,000-7,000 Da, and a mass step of 0.1 Da.

### StdGH enzymatic assay

Lyophilized StdA was dissolved in buffer (50 mM HEPES pH 8, 150 mM KCl) to a final concentration of 1 mM and then frozen in individual aliquots at -80 °C until used. For the first step of the biosynthesis pathway (thioamidation), aliquots of StdA and purified StdGH were thawed on ice. The 200 µL reaction mixture contained 100 µM StdA, 20 µM StdGH, 50 mM HEPES pH 8, 150 mM KCl, 5 mM Na_2_S, 5 mM ATP, 5 mM MgCl_2_, and 1 mM DTT. The reaction was incubated at room temperature overnight on the bench. The next day, the reaction mixture was heated to 70 °C for 10 min then centrifuged at 8000 × g to remove StdGH. The supernatant containing the StdGH-modified StdA product was dialyzed into Buffer C using a 2K MWCO Slide-A-Lyzer™ Dialysis Cassette, following the manufacturer’s instructions to remove the excess Na_2_S, ATP, MgCl_2,_ and DTT. The samples were then analyzed directly using intact mass spectrometry described above.

### StdF enzymatic assay

StdGH-modified StdA was used as the substrate for the StdF enzyme assay. The 200 µL reaction mixture contained 100 µM StdGH-modified StdA, 20 µM StdF, 50 mM HEPES pH 8, 150 mM KCl, 1 mM carbamoyl phosphate, 5 mM ATP, and 5 mM MgCl_2_. The reaction was incubated at room temperature overnight on the bench. The following day, the reaction mixture was heated to 70 °C for 10 min and then centrifuged at 8000 × g to remove StdF. The supernatant containing the StdGHF-modified StdA product was then analyzed directly using intact mass spectrometry described above.

### StdDE enzymatic assay

The StdDE enzyme assay was performed in the presence of StdF to immediately transfer the cleaved, carbamoylated, and thioamidated substrate to StdDE. StdDE was prepared with a 5-fold molar excess of ascorbate and ammonium iron(II) sulfate hexahydrate on the benchtop before the reaction. The 200 µL reaction mixture contained 100 µM StdGH-modified StdA, 20 µM StdF, 20 µM StdDE, 50 mM HEPES pH 8, 150 mM KCl, 1 mM carbamoyl phosphate, 5 mM ATP, 5 mM MgCl_2_, 100 µM ascorbate, and 100 µM ammonium iron(II) sulfate hexahydrate. The reaction was incubated on the bench at room temperature overnight. The following day, the reaction mixture was heated to 70 °C for 10 min, then centrifuged at 8000 × g to remove StdF and StdDE. The supernatant containing StdGHFDE-modified StdA was then analyzed using intact mass spectrometry described above.

### Isolation of unmodified and modified core-containing fragments

To isolate unmodified and modified core-containing fragments, proteolysis was performed for 45 min at 37 °C using ArgC-Ultra at an enzyme-to-substrate ratio of 1:500. The reactions were then quenched by adding formic acid to a final concentration of 0.1%. HPLC purification was performed to isolate the fragments on an Agilent 1100 system in the Northwestern Keck Biophysics Facility. The column used was a Phenomenex Aeris Peptide XB-C18 LC column (Part no. 00G-4632-N0; particle size: 5 µm; dimensions: 250 x 10 mm; pore size: 100 Å) coupled with a SecurityGuard SemiPrep Cartridge core-shell C18-Peptide (Part no. AJ0-9317; dimensions: 10 x 10 mm). The mobile phase consisted of solvent A, which contained water with 0.5% acetic acid, and solvent B, with acetonitrile and 0.5% acetic acid. The column was equilibrated with 10 column volumes of 99% solvent A and 1% solvent B. An injection volume of 500 µL was used. Separation was performed over a gradient of 1-30% solvent B at a flow rate of 2 mL/min for 30 min. The column was then washed with 95% solvent B for 5 min at 2.5 mL/min, followed by re-equilibration with 1% solvent B for 10 min at 2 mL/min. The products were freeze-dried to remove the acetonitrile/water/0.5% acetic acid solution. Retention times of the Arg-C Ultra digested unmodified StdA, StdAGH-modified StdA, StdAGHF-modified StdA, and StdAGHFDE-modified StdA are listed in Table S11.

### High-resolution tandem mass spectrometry (HR-MS/MS)

HR-MS/MS data collection and analysis were performed in the Northwestern Proteomics Core Facility. Briefly, the raw data for the Arg-C Ultra digested samples were acquired on an Orbitrap Exploris 240 (Lucy) using data-dependent acquisition (DDA) mode. The data were processed in FragPipe against the reference peptide sequence with a universal list of common contaminants included.

### Large-scale enzymatic reactions to obtain Arg-C Ultra digested unmodified ^15^N/^13^C-Asn13-Lys14 StdA

To obtain the appropriate sample amount for NMR, the unmodified ^13^C and ^15^N labeled StdA (20 mg) was dissolved in 5 mL of buffer consisting of 50 mM HEPES pH 8, 150 mM KCl. Arg-C Ultra digestion was performed as described above. The digestion was then sent to the Peptide Synthesis Core Facility of the Center of Regenerative Nanomedicine at Northwestern University for purification of the Arg-C Ultra digested unmodified ^15^N/^13^C-Asn13-Lys14 StdA.

### Large-scale enzymatic reactions and purification to obtain Arg-C Ultra digested StdGHFDE-modified ^15^N/^13^C-Asn13-Lys14 StdA

To obtain the appropriate sample amount for NMR, the unmodified ^13^C and ^15^N labeled StdA (65 mg) was dissolved in 10 mL of buffer consisting of 50 mM HEPES pH 8, 150 mM KCl, 20 mM ATP, 20 mM MgCl_2_, and 20 mM Na_2_S. The StdGH enzyme complex was then added to the mixture at a final concentration of 100 μM. The reaction was conducted overnight on the benchtop at room temperature. The next day, the reaction was aliquoted into 1.5 mL microcentrifuge tubes and heated at 70 °C for 10 min to remove StdGH. The remaining StdGH-modified ^15^N/^13^C-Asn13-Lys14 StdA was dialyzed into Buffer C using a 2K MWCO Slide-A-Lyzer™ Dialysis Cassette, following the manufacturer’s instructions to remove the excess Na_2_S, ATP, MgCl_2,_ and DTT. The 20 mL reaction mixture for the combined StdF and StdDE reaction contained 500 μM StdGH-modified ^15^N/^13^C-Asn13-Lys14 StdA, 100 μM StdF, 100 μM StdDE, 50 mM HEPES pH 8, 150 mM KCl, 1 mM carbamoyl phosphate, 5 mM ATP, 5 mM MgCl_2_, 500 µM ascorbate, and 500 µM ammonium iron(II) sulfate hexahydrate. The reaction was left on the benchtop at room temperature overnight. The next day, aliquots were heated at 70 °C to remove the enzymes and centrifuged at 8000 × g to obtain the supernatant. Arg-C Ultra digestion was performed as described above. The digestion was then sent to the Peptide Synthesis Core Facility of the Center of Regenerative Nanomedicine at Northwestern University for purification of the Arg-C Ultra digested StdGHFDE-modified ^15^N/^13^C-Asn13-Lys14 StdA.

### NMR data acquisition and analysis

All spectra were collected at 25 °C on a Bruker HCFN 600 MHz spectrometer at Northwestern’s IMSERC facility, equipped with a triple channel, QCI cryo probe with Z gradients. Assignment experiments were carried out at natural ^13^C/^15^N abundance on the modified StdA peptide at a concentration of 200 µM in both H_2_O and D_2_O using uniform spectral sampling. The TOCSY experiments were collected using States-TPPI acquisition with sweep widths of 8197 and 7198 Hz in the F2 and F1 dimensions with respective resolutions of 4.0 and 28.1 Hz in both solvents. Homonuclear Hartman-Hahn transfers were accomplished with DIPSI2 and excitation sculpting was achieved with gradients. ROESY experiments were acquired in States-TPPI mode with sweep widths of 7143 Hz in both F2 and F1 and resolutions of 3.5 and 27.9 Hz in F2 and F1 in both solvents. A cw spinlock was used with a mixing time of 200 ms. A D_2_O-COSY was collected using QF acquisition with sweep widths of 7143 and 7198 Hz in F2 and F1 with respective resolutions of 3.5 and 28.1 Hz. Gradient pulses were used for selection. A multiplicity-edited, D_2_O ^13^C-HSQC was collected using echo-antiecho acquisition with sweep widths of 8197 and 24887 Hz in F2 and F1 with respective resolutions of 8.0 and 388.9 Hz. Magnetization transfer was accomplished using double INPET transfers optimized for sensitivity enhancement and decoupling was used during acquisition.

The HNCO, low pH ^15^N-HSQCs, and comparative ^13^C-HSQCs were collected on unmodified and StdGHFDE-modified U-[^13^C,^15^N]-Asn13-Lys14-StdA at 1500 and 700 μM, respectively, in 20 mM sodium phosphate pH 6.8, 50 mM NaCl, 1% DSS, 10% D_2_O. The ^15^N-HSQCs were non-uniformly sampled, phase-sensitive, double INPET transfer-based sequences with Watergate solvent decoupling. They had ^15^N offsets of 105 ppm with sweep widths of 6849 and 3039 Hz in ^1^H and ^15^N, leading to resolutions of 6.7 and 23.7 Hz respectively. The HNCOs were acquired as pseudo-2D experiments with one time domain point for the ^15^N dimension, sweep widths of 8197 and 4525 Hz in ^1^H and ^13^C giving resolutions of 8.0 and 70.7 Hz respectively. The ^13^C-HSQCs were non-uniformly sampled, multiplicity-edited, echo-antiecho acquired, double INPET transfer based sequences. A ^13^C offset of 40 ppm was used with sweep widths of 8197 and 12066 Hz in F2 and F1 with respective resolutions of 8 and 94.3 Hz.

## ASSOCIATED CONTENT

### Supporting Information

Supplementary Tables S1-S11, Supplementary Figures S1-S14

## Notes

The Authors declare no competing financial interests.

## ACKNOWLEDGMENTS

StdA peptide synthesis was performed at the Peptide Synthesis Core Facility of the Center for Regenerative Nanomedicine at Northwestern University. ICP-MS analysis was performed at the Quantitative Bio-element Imaging Center at Northwestern University, supported by NASA Ames Research Center Grant NNA04CC36G. HPLC purification was conducted in the Northwestern Keck Biophysics Facility, a shared resource of the Robert H. Lurie Comprehensive Cancer Center of Northwestern University supported in part by the NCI CCSG P30 CA060553. Proteomics services were performed by the Northwestern Proteomics Core Facility (RRID:SCR_017945), generously supported by NCI CCSG P30 CA060553 awarded to the Robert H Lurie Comprehensive Cancer Center, instrumentation award (S10OD025194) from NIH Office of the Director, and the National Resource for Translational and Developmental Proteomics supported by P41 GM108569. This work made use of the IMSERC (RRID:SCR_017874) NMR facility at Northwestern University, which has received support from the Soft and Hybrid Nanotechnology Experimental (SHyNE) Resource (NSF ECCS-2025633), NIH 1S10OD012016-01 / 1S10RR019071-01A1, and Northwestern University.

