## Supporting Information for "Bis-hydroxylation of Homocitrulline Catalyzed by a Multinuclear Nonheme Iron-Dependent Oxidative Enzyme during RiPP Biosynthesis"

Figures S1-S14
Tables S1-S10
DNA and amino acid sequences

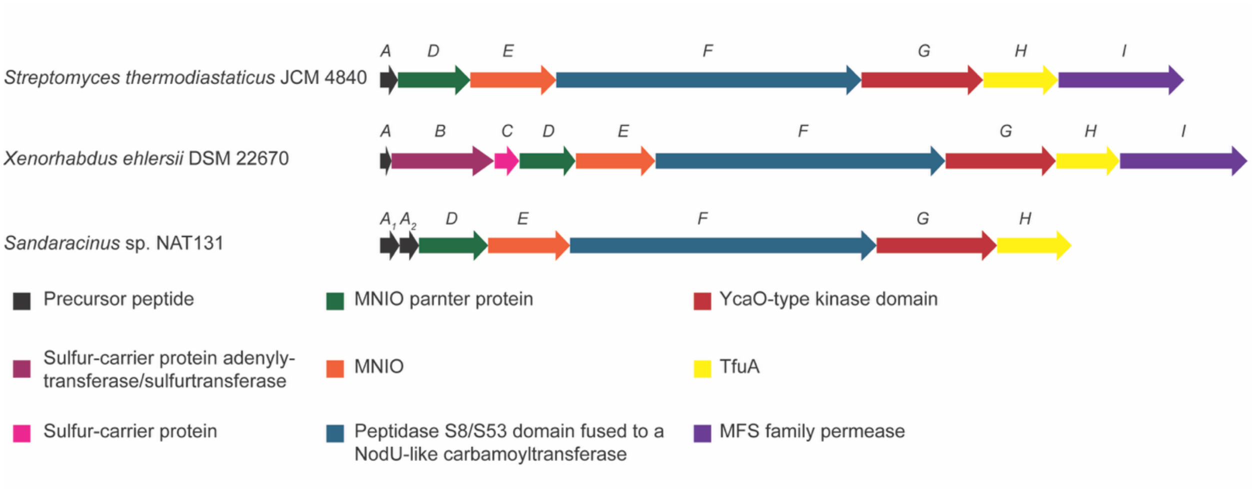


**Figure S1.** Representative BGCs from Actinomycetota, Pseudomonadota, and Myxococcota phyla.

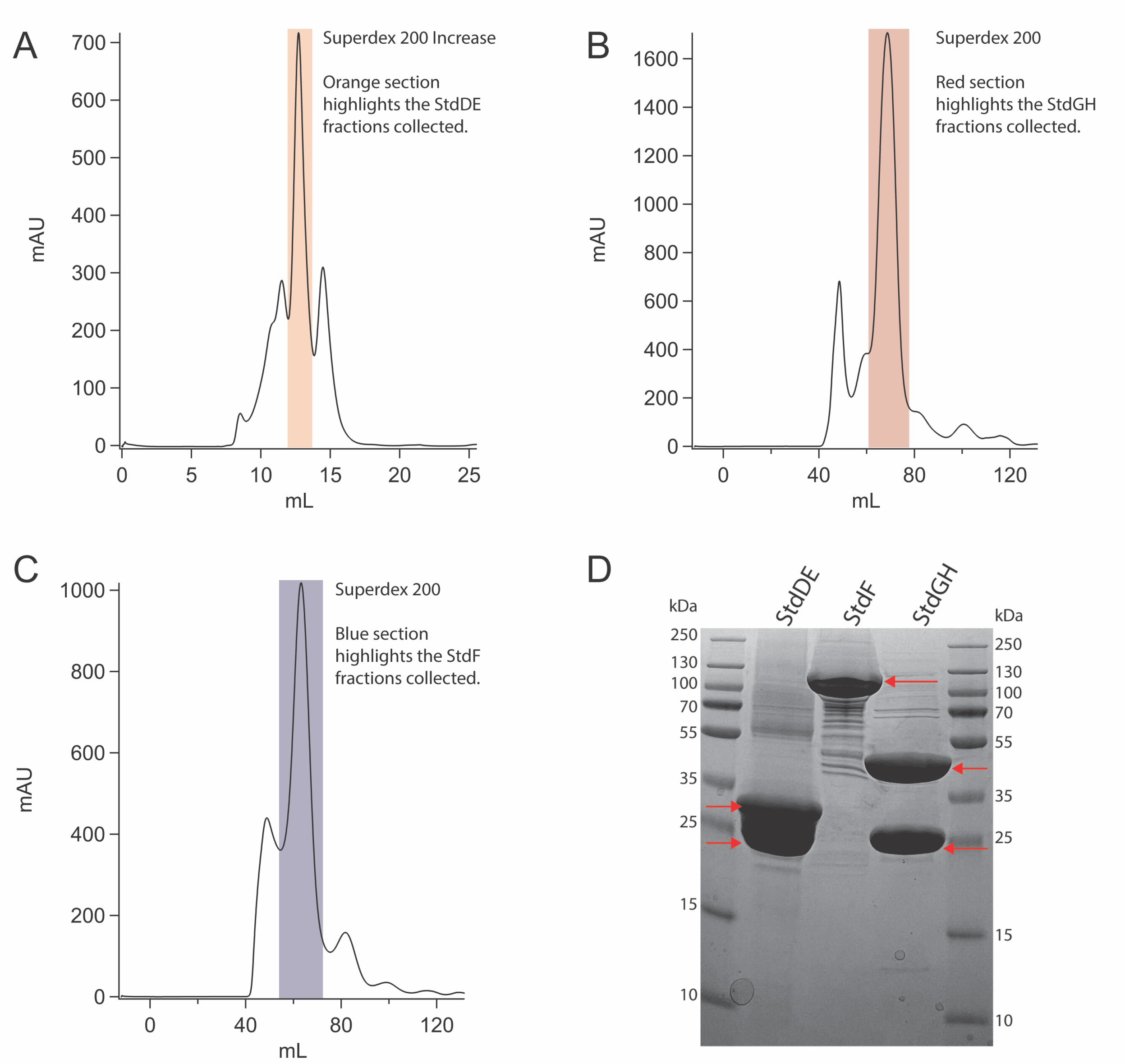


**Figure S2.** Purification of StdGH, StdDE, and StdF. SEC trace of (A) StdDE, (B) StdGH, and (C) StdF. (D) SDS-PAGE analysis of purified StdD (28.9 kDa) - StdE (32.7 kDa) complex, StdF (114.5 kDa), and StdG (45.4 kDa) - StdH (28.7 kDa) complex.

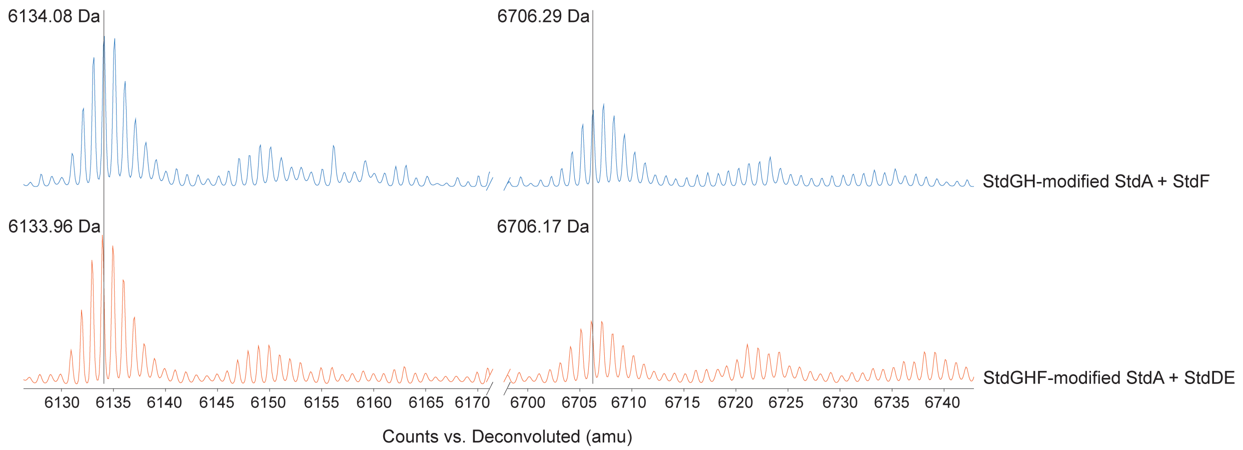


**Figure S3.** LC-TOF-MS of StdGHF-modified peptide reacted with StdDE, ascorbate, and ammonium iron(II) sulfate hexahydrate.

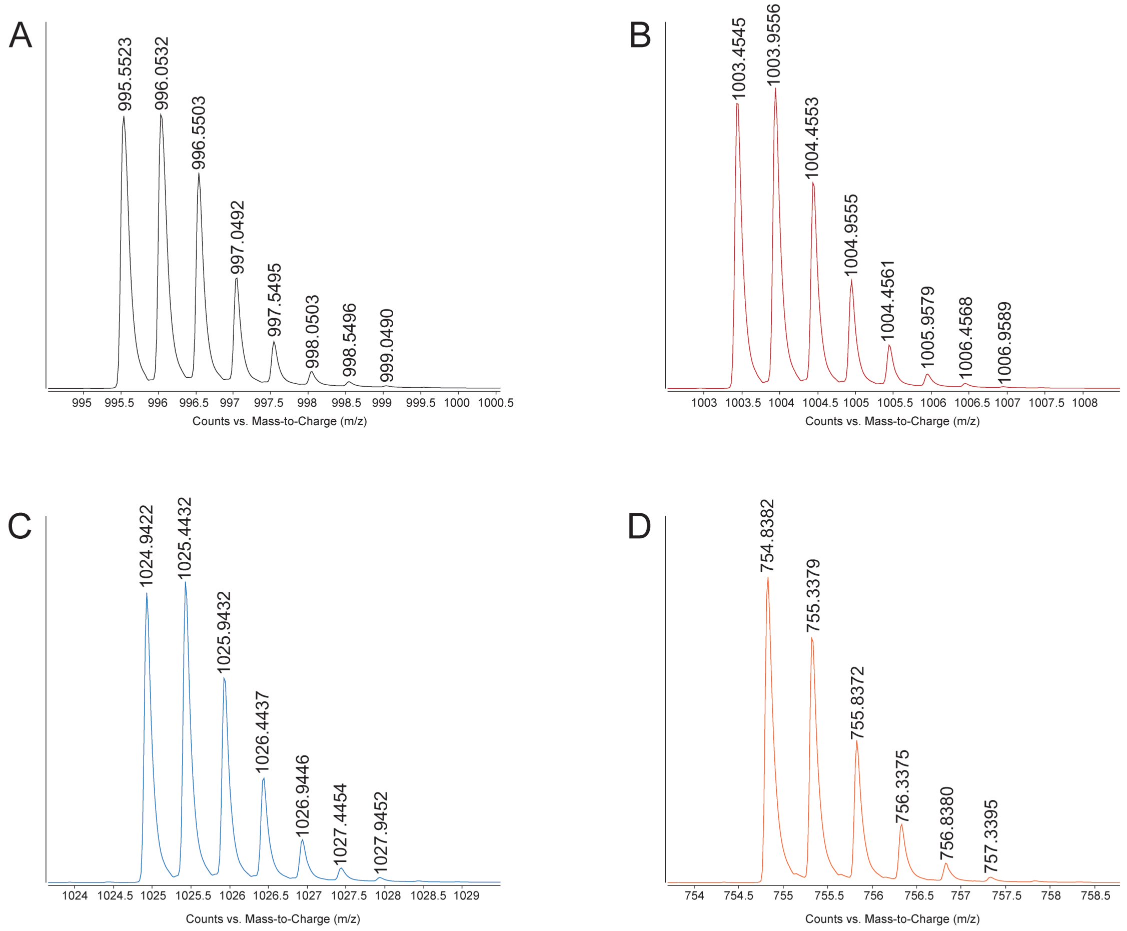


**Figure S4.** LC-ESI focusing on the $[M+2H]^{2+}$ ion of the Arg-C Ultra fragments of (A) unmodified StdA, (B) StdGH-modified StdA, (C) StdGHF-modified StdA, and (D) StdGHFDE-modified StdA peptides.

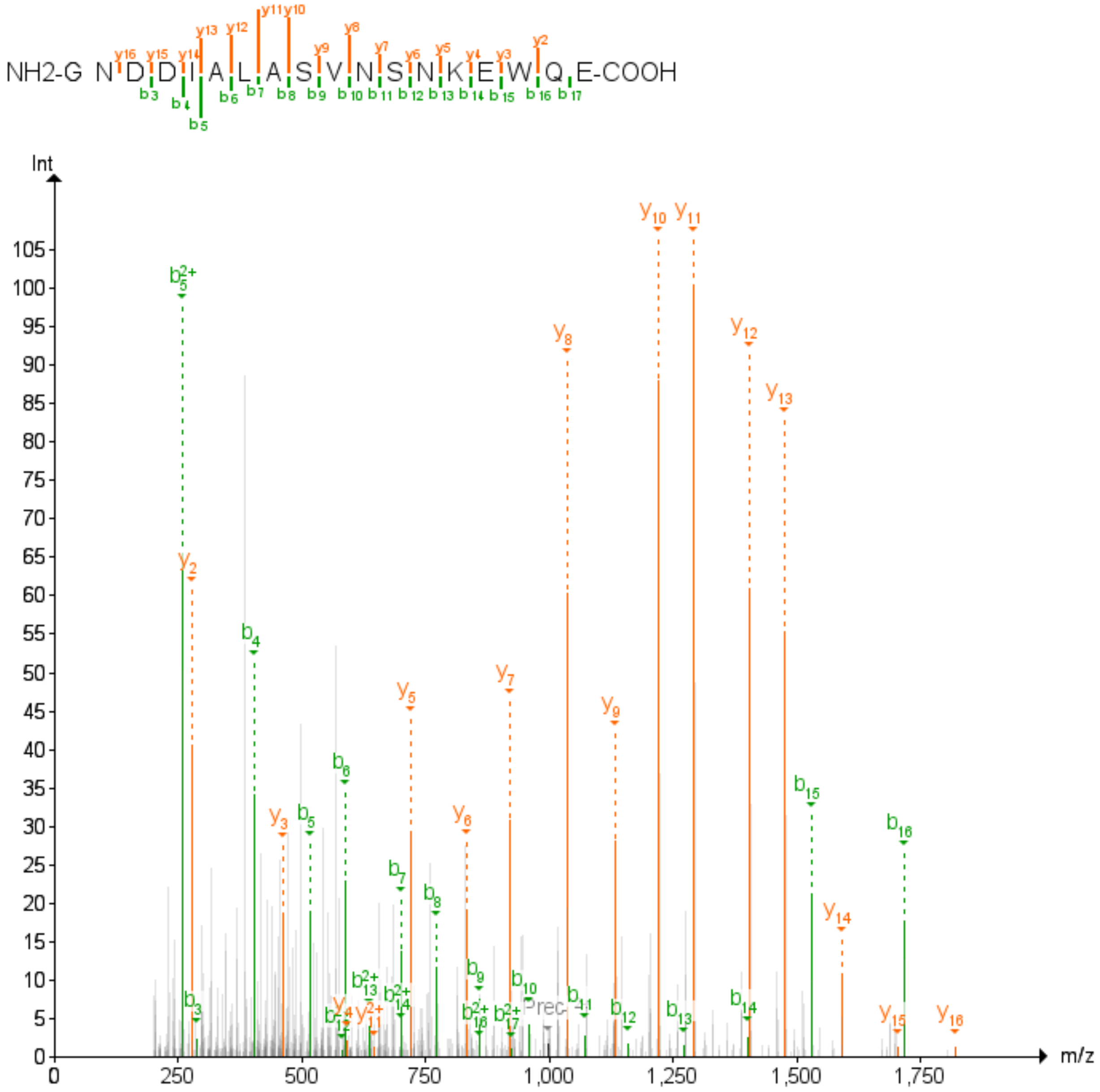


**Figure S5.** HR-MS/MS spectra of Arg-C Ultra digested unmodified StdA.

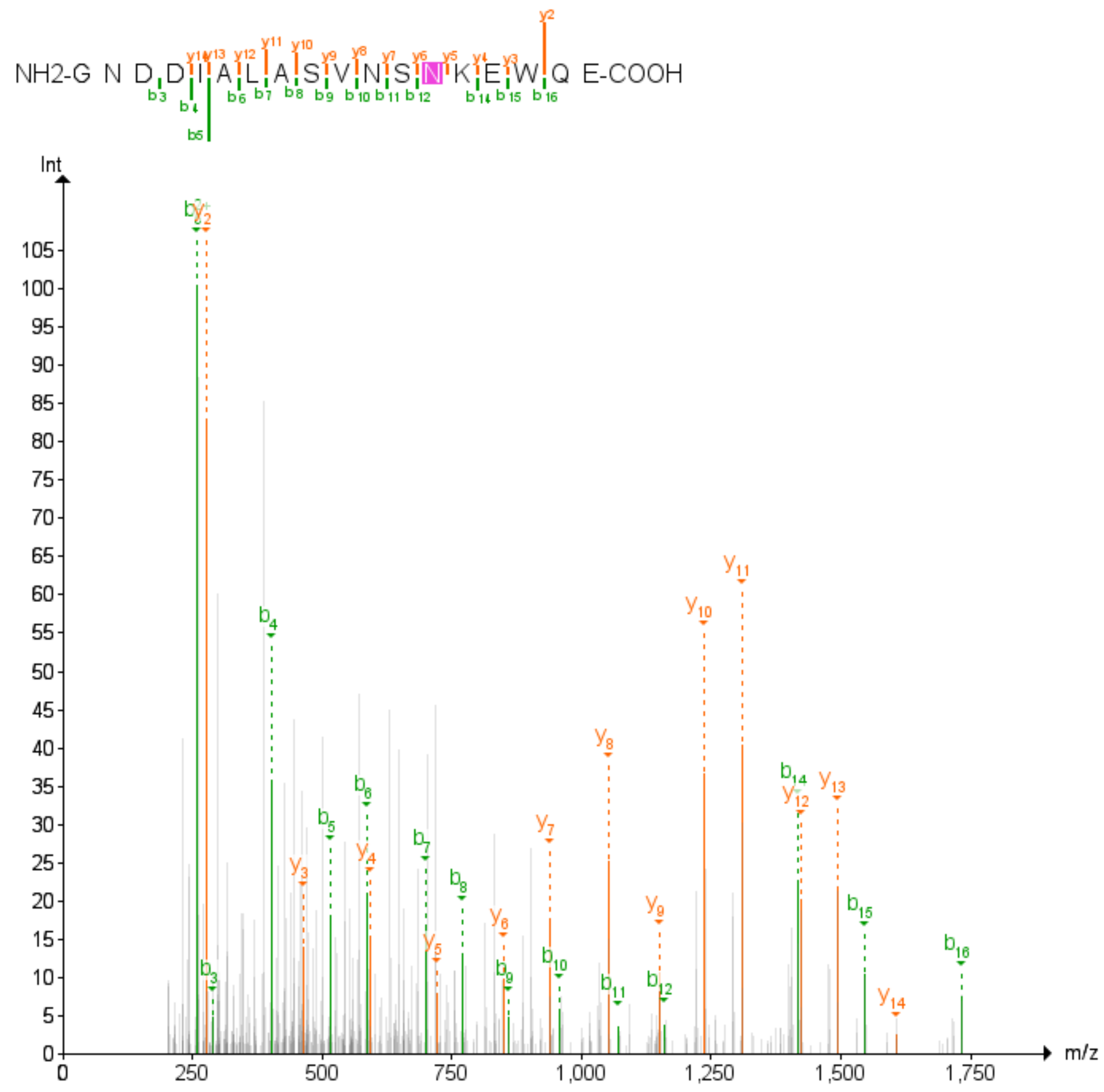


**Figure S6.** HR-MS/MS spectra of Arg-C Ultra digested StdA peptide following the in vitro reaction with StdGH in the presence of ATP and $Na_2S$.

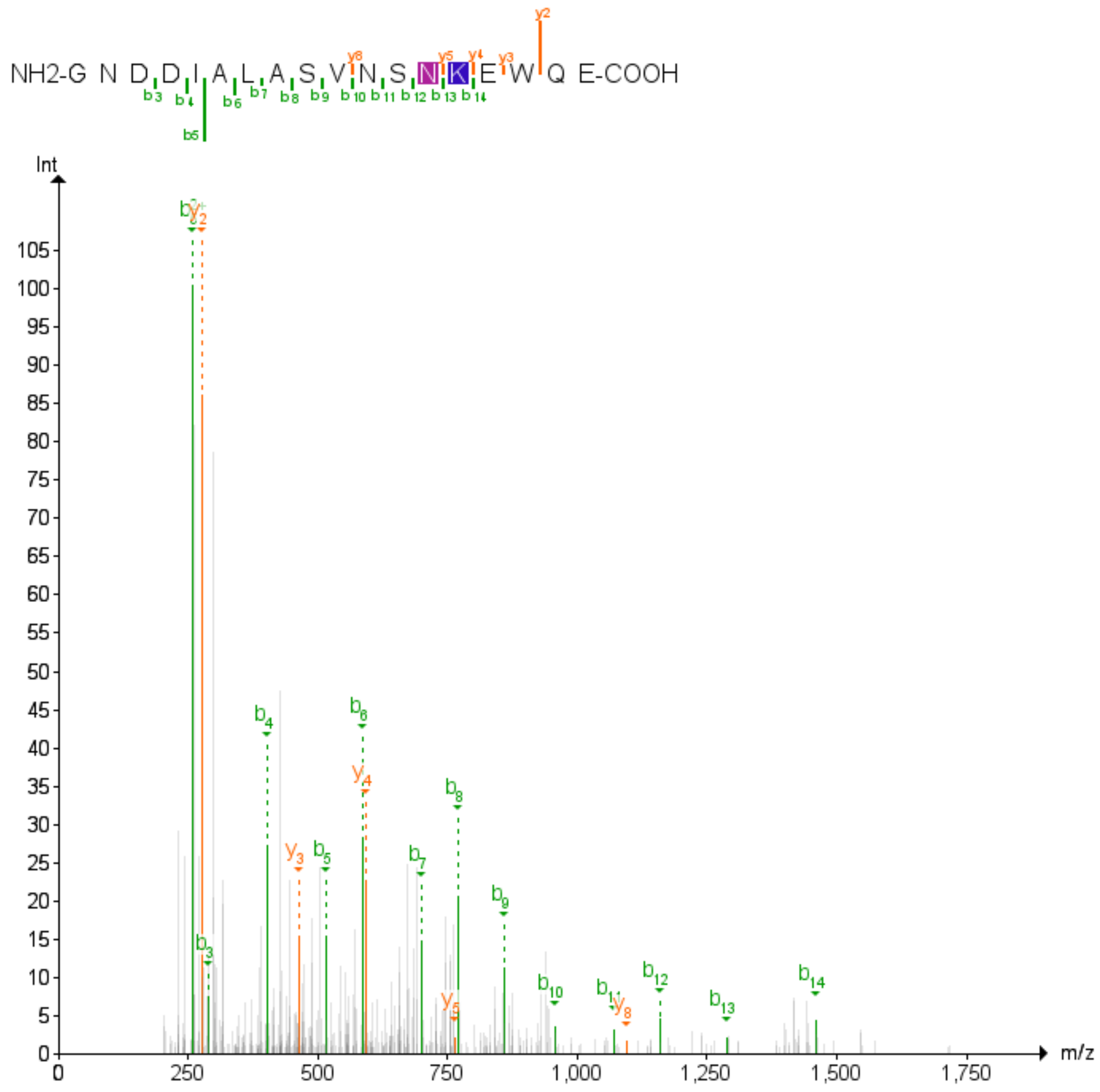


**Figure S7.** HR-MS/MS spectra for Arg-C Ultra digested for StdGH-modified StdA peptide following the in vitro reaction with StdF in the presence of carbamoyl phosphate and ATP. This species represents the full-length peptide.

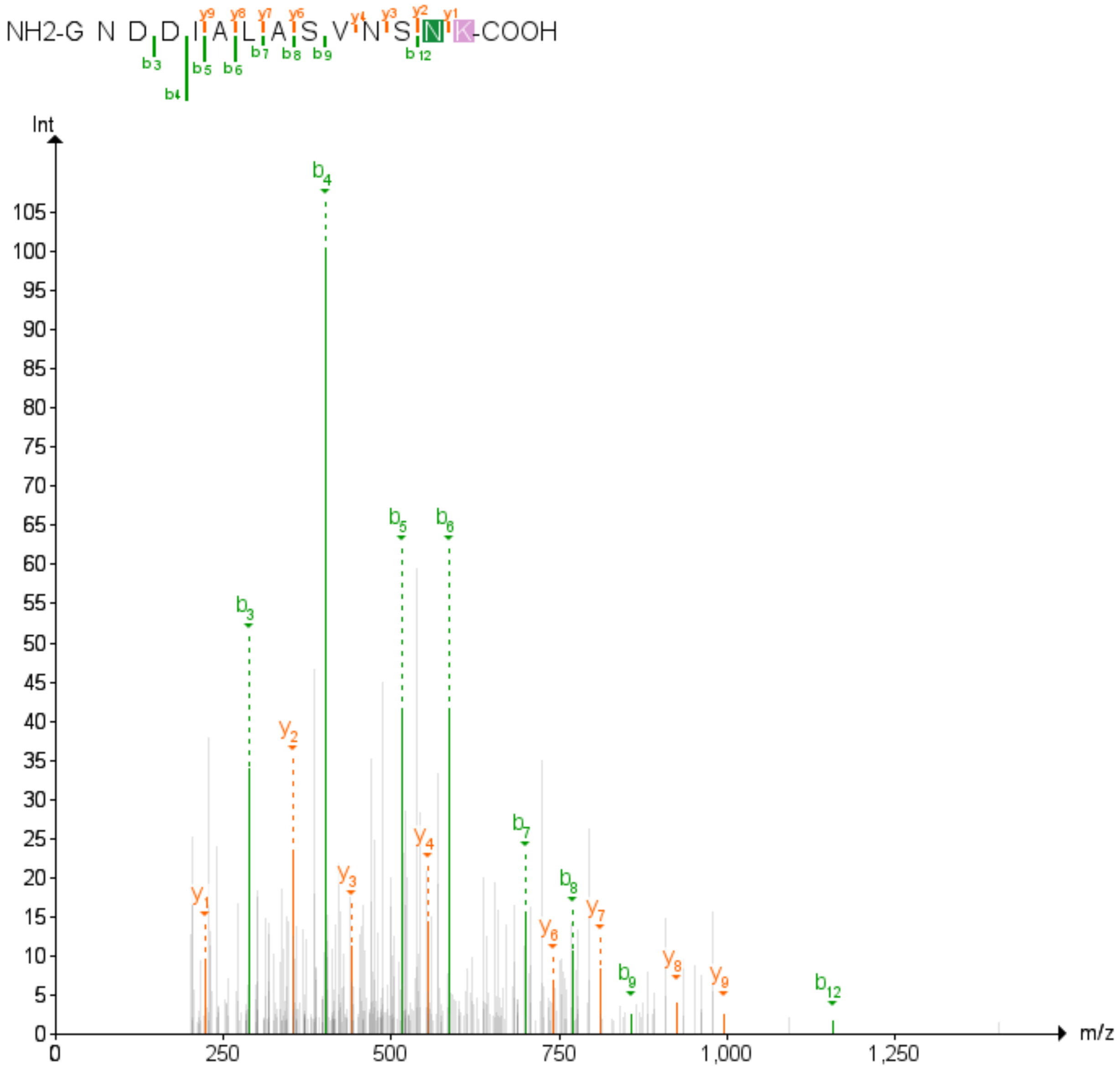


**Figure S8.** HR-MS/MS spectra for Arg-C Ultra digested for cleaved StdGH-modified StdA peptide following the in vitro reaction with StdF in the presence of carbamoyl phosphate and ATP, and StdDE in the presence of iron and ascorbate.

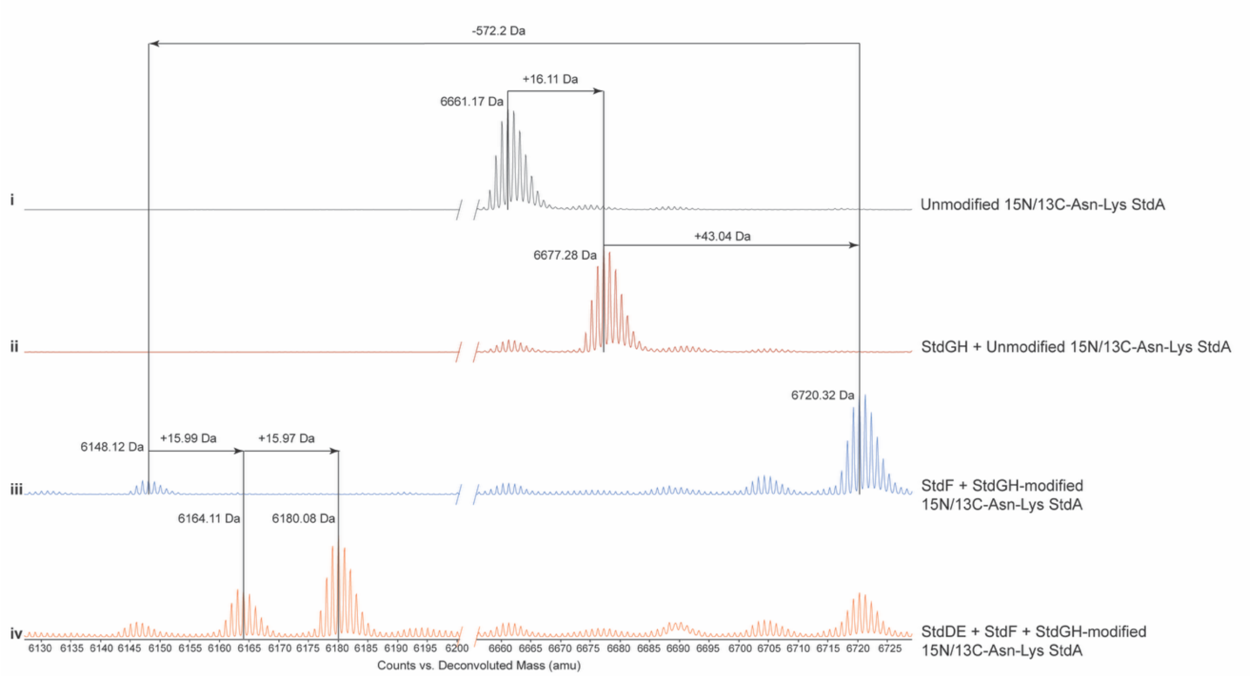


**Figure S9.** In vitro reactions of StdGH, StdF, and StdDE with $^{13}C/^{15}N$-Asn56-Lys57-StdA monitored by LC-TOF-MS.

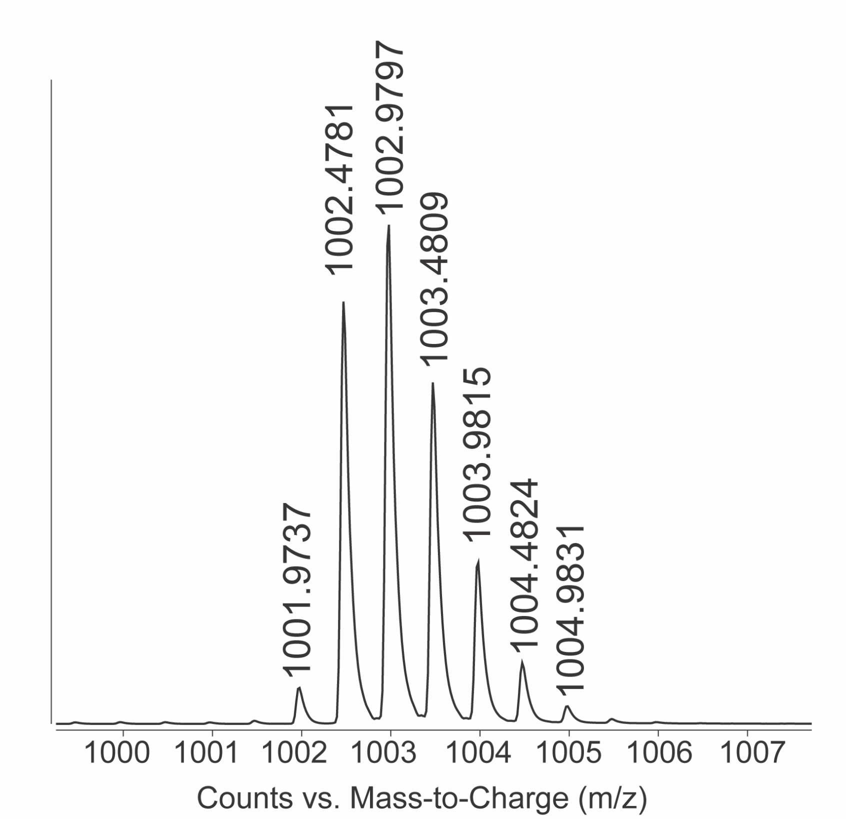


**Figure S10.** LC-ESI focusing on the $[M+2H]^{2+}$ ion of Arg-C Ultra fragment derived from the 6661.17 Da unmodified $^{15}N/^{13}C$-Asn13-Lys14-StdA.

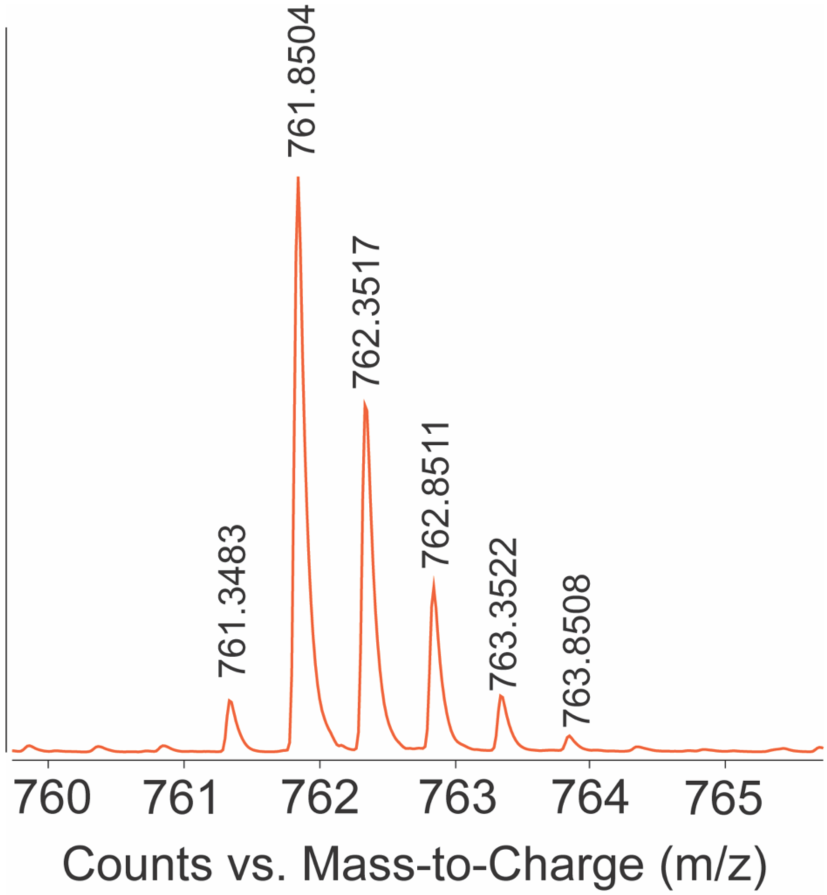


**Figure S11.** LC-ESI focusing on the $[M+2H]^{2+}$ ion of Arg-C Ultra fragment derived from the 6180.08 Da StdGHFDE-modified $^{15}N/^{13}C$-Asn13-Lys14-StdA.

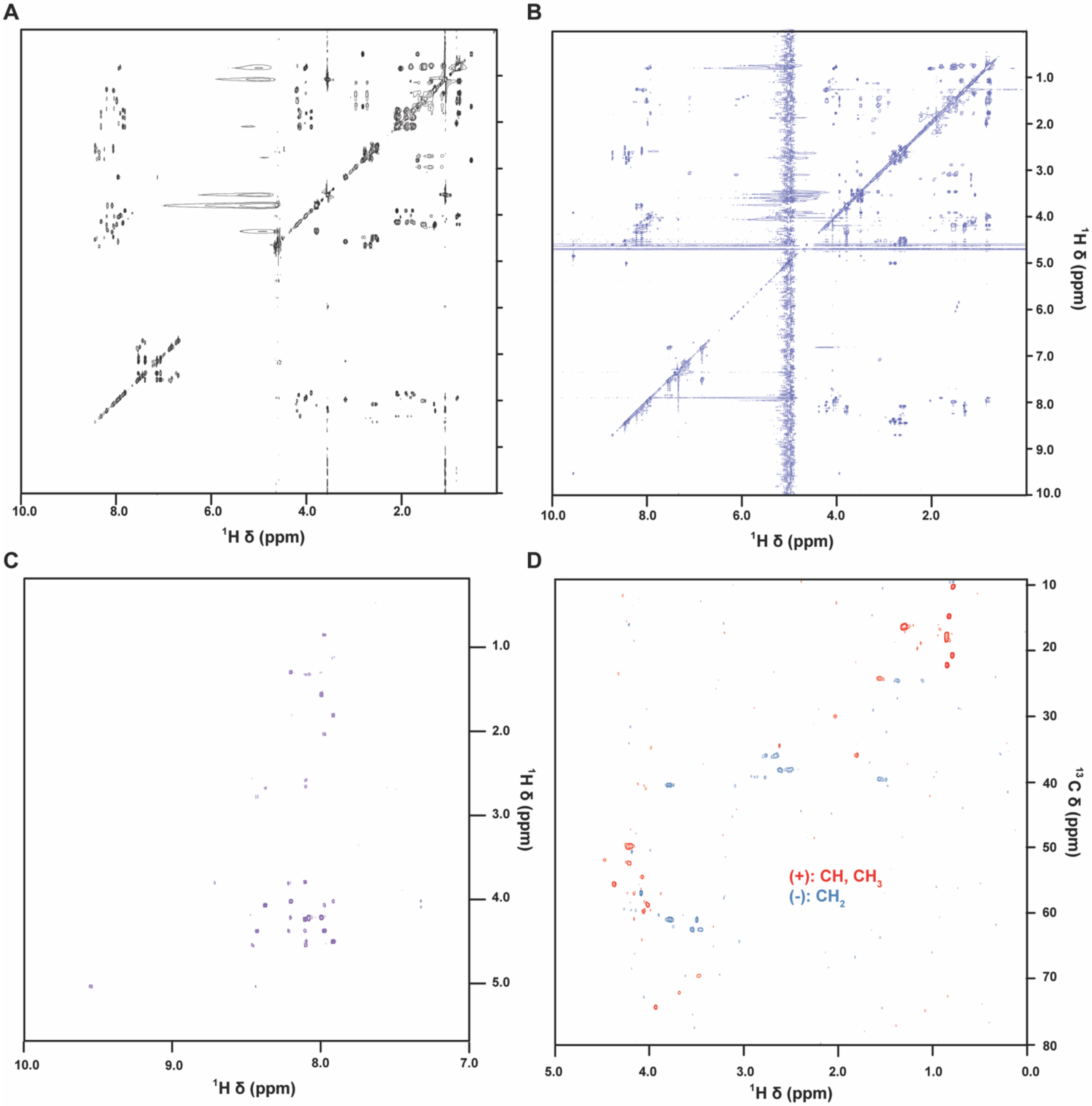


**Figure S12.** Assignment of Arg-C Ultra digested StdGHFDE-modified $^{15}N/^{13}C$-Asn13-Lys14-StdA. (A) TOCSY spectrum of unmodified $^{15}N/^{13}C$-Asn13-Lys14-StdA. (B) TOCSY spectrum of modified StdA. (C) HN- contacts in StdGHFDE-modified $^{15}N/^{13}C$-Asn13-Lys14-StdA ROESY spectrum with 200 ms mixing time. (D) Multiplicity-edited $^{13}C$-HSQC spectrum.

**[1]GNDDIALAS[10]VNSN*K***

**[1]Gly**
$C^{\alpha}$: 40.5 $H^{\alpha 1}$: 3.68 $H^{\alpha 2}$: 3.65

**[2]Asn**
$H^{N}$: 8.59 $H^{\alpha}$: 4.50 $H^{\beta 1}$: 2.64
$C^{\alpha}$: 54.0 $C^{\beta}$: 35.9 $H^{\beta 2}$: 2.53

**[3]Asp**
$H^{N}$: 8.33 $H^{\alpha}$: 4.40 $H^{\beta 1}$: 2.50
$C^{\alpha}$: 51.7 $C^{\beta}$: 38.3 $H^{\beta 2}$: 2.37

**[4]Asp**
$H^{N}$: 7.97 $H^{\alpha}$: 4.35 $H^{\beta 1}$: 2.48
$C^{\alpha}$: 51.9 $C^{\beta}$: 38.0 $H^{\beta 2}$: 2.41

**[5]Ile**
$H^{N}$: 7.79 $C^{\gamma 1}$: 24.6 $H^{\gamma 21}$: 0.700
$C^{\alpha}$: 58.7 $H^{\gamma 11}$: 1.243 $H^{\gamma 22}$: 0.689
$H^{\alpha}$: 3.89 $H^{\gamma 12}$: 0.981 $C^{\delta}$: 10.2
$C^{\beta}$: 35.9 $C^{\gamma 2}$: 14.7 $H^{\delta}$: 0.656
$H^{\beta}$: 1.67

**[6]Ala**
$H^{N}$: 7.95 $H^{\alpha}$: 4.10 $H^{\beta 1}$: 1.20
$C^{\alpha}$: 52.4 $C^{\beta}$: 16.4 $H^{\beta 2}$: 1.19

**[7]Leu**
$H^{N}$: 7.87 $H^{\beta 2}$: 1.36 $H^{\delta 12}$: 0.714
$C^{\alpha}$: 52.4 $C^{\gamma}$: 24.2 $C^{\delta 2}$: 20.7
$H^{\alpha}$: 4.08 $H^{\gamma}$: 1.44 $H^{\delta 21}$: 0.666
$C^{\beta}$: 39.6 $C^{\delta 1}$: 22.2 $H^{\delta 22}$: 0.654
$H^{\beta 1}$: 1.40 $H^{\delta 11}$: 0.722

**[8]Ala**
$H^{N}$: 8.07 $H^{\alpha}$: 4.09 $H^{\beta 1}$: 1.17
$C^{\alpha}$: 49.8 $C^{\beta}$: 16.2 $H^{\beta 2}$: 1.16

**[9]Ser**
$H^{N}$: 7.98 $H^{\alpha}$: 4.24 $H^{\beta}$: 3.66
$C^{\alpha}$: 55.4 $C^{\beta}$: 60.928

**[10]Val**
$H^{N}$: 7.85 $H^{\beta}$: 1.90 $C^{\gamma 2}$: 17.6
$C^{\alpha}$: 59.7 $C^{\gamma 1}$: 18.3 $H^{\gamma 21}$: 0.732
$H^{\alpha}$: 3.93 $H^{\gamma 11}$: 0.728 $H^{\gamma 22}$: 0.717
$C^{\beta}$: 30.0 $H^{\gamma 12}$: 0.714

**[11]Asn**
$H^{N}$: 8.25 $H^{\alpha}$: 4.58 $H^{\beta 1}$: 2.67
$C^{\alpha}$: 55.1 $C^{\beta}$: 36.1 $H^{\beta 2}$: 2.55

**[12]Ser**
$H^{N}$: 8.04 $H^{\alpha}$: 4.26 $H^{\beta}$: 3.66
$C^{\alpha}$: 55.6 $C^{\beta}$: 60.930

**[13]Modified-Asn**
$H^{N}$: 8.33 $H^{\alpha}$: 4.90 $C^{\gamma}$: 174.3
N: 125.1 $C^{\beta}$: 39.3 $N^{\delta}$: 113.4
$C^{S}$: 171.5 $H^{\beta 1}$: 2.76 $H^{\delta 1}$: 7.40
$C^{\alpha}$: 56.1 $H^{\beta 2}$: 2.65 $H^{\delta 2}$: 6.75

**[13]Unmodified-Asn**
$H^{N}$: 8.26 $H^{\alpha}$: 4.34 $C^{\gamma}$: 174.3
N: 121.3 $C^{\beta}$: 38.2 $N^{\delta}$: 113.4
$C^{O}$: 172.4 $H^{\beta 1}$: 2.59 $H^{\delta 1}$: 7.40
$C^{\alpha}$: 52.8 $H^{\beta 2}$: 2.50 $H^{\delta 2}$: 6.75

**[14]Modified-Lys**
$H^{N}$: 9.43 $C^{\gamma}$: 69.5 $H^{\varepsilon}$: 3.00
N: 112.2 $H^{\gamma}$: 3.35 $N^{\zeta}$: 87.0
$C^{\alpha}$: 62.5 $C^{\delta}$: 32.5 $H^{\zeta}$: 6.00
$H^{\alpha}$: 4.74 $H^{\delta 1}$: 1.52 $C^{\eta}$: 171.4
$C^{\beta}$: 74.3 $H^{\delta 2}$: 1.36 $N^{\theta}$: 117.5
$H^{\beta}$: 3.81 $C^{\varepsilon}$: 45.3 $H^{\theta}$: 7.65

**[14]Unmodified-Lys**
$H^{N}$: 8.06 $H^{\beta}$: 1.51 $C^{\delta}$: 21.8
N: 112.2 $C^{\gamma}$: 30.2 $H^{\delta}$: 121.1
$C^{\alpha}$: 53.7 $H^{\gamma 1}$: 1.62 $C^{\varepsilon}$: 39.4
$H^{\alpha}$: 4.13 $H^{\gamma 2}$: 1.58 $H^{\varepsilon}$: 2.83
$C^{\beta}$: 26.3

**Figure S13.** Arg-C Ultra digested StdGHFDE-modified $^{15}N/^{13}C$-Asn13-Lys14-StdA chemical shift list. Complete chemical shift assignments for the StdGHFDE-modified $^{15}N/^{13}C$-Asn13-Lys14-StdA and unmodified $^{15}N/^{13}C$-Asn13-Lys14-StdA Asn13 and Lys14 residues. Amino acid sequence shown at top with sites of modification indicated by an asterisk.

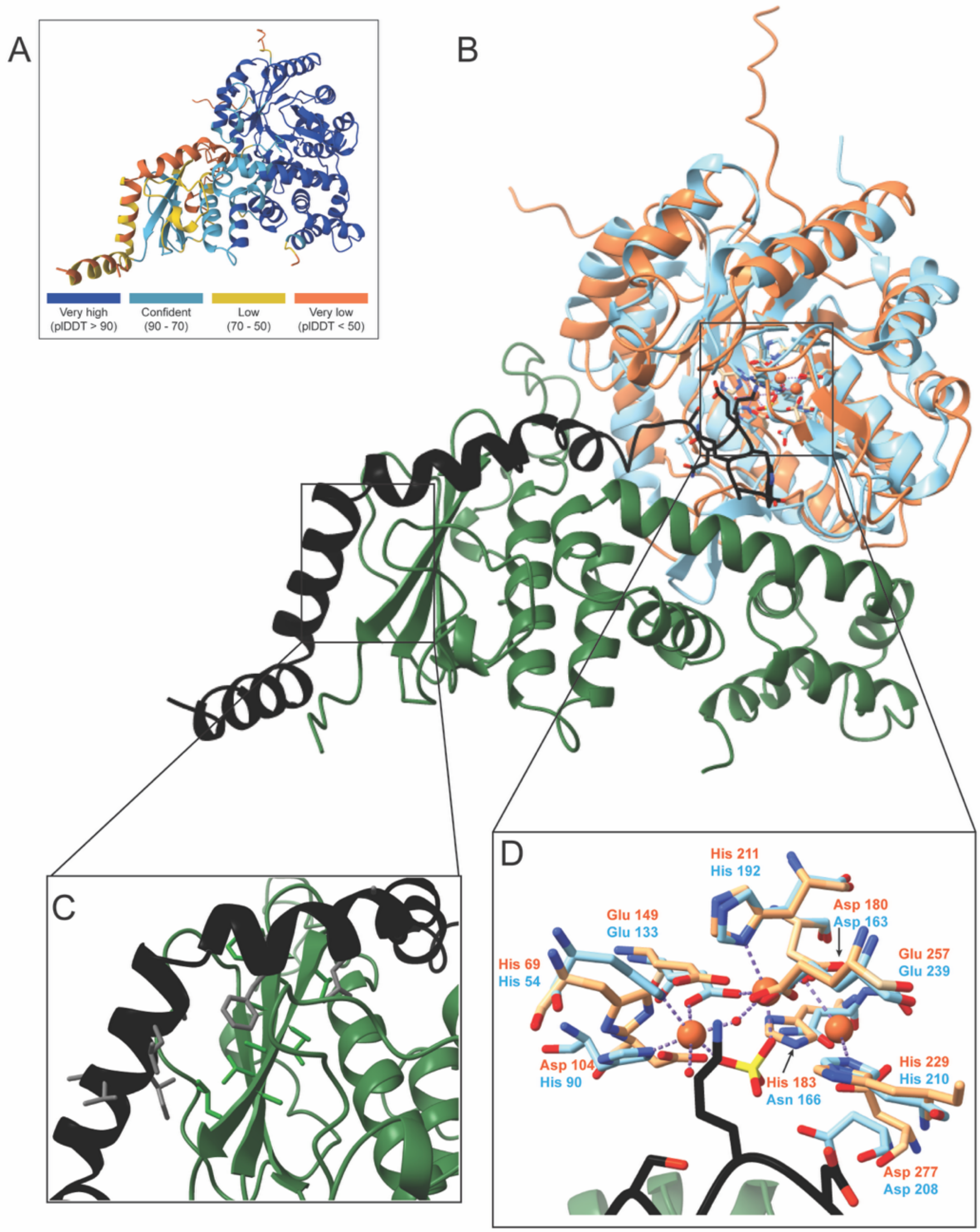


**Figure S14. AlphaFold 3 model of StdDE in complex with cleaved StdA.** (A) Confidence metrics for the AlphaFold 3-predicted structure. Per-residue pLDDT scores are shown using the color scale indicated in the legend, with higher values indicating higher confidence. (B) AlphaFold 3-predicted model of the StdDE–StdA complex showing StdA (black), StdD (green), and StdE (orange). StdE is superimposed with the crystal structure of MbnB from *Methylosinus trichosporium* OB3b (blue; PDB ID: 7TCR). (C) Close-up view of the predicted RRE site in StdD (green), highlighting hydrophobic residues interacting with hydrophobic residues in the StdA leader peptide (black). (D) Conserved residues in StdE are predicted to coordinate Fe ions and modify StdA Lys57, based on comparison with the MbnB crystal structure.

**Table S1.** IMG Gene ID, gene names, and predicted functions of genes in the *std* cluster.

| Gene ID | Gene name | Predicted function |
|---|---|---|
| 8109937978 | *stdA* | Precursor peptide |
| 8109937979 | *stdD* | MNIO partner protein |
| 8109937980 | *stdE* | MNIO |
| 8109937981 | *stdF* | Peptidase S8/S53 domain fused to a NodU-like carbamoyltransferase |
| 8109937982 | *stdG* | YcaO-type kinase domain |
| 8109937983 | *stdH* | TfuA |
| 8109937984 | *stdI* | MFS family permease |

**Table S2**. Prevalence of StdE in microbial genomes. IMG Gene ID and strains are provided. Entries are in order from the highest sequence identity (%ID) to the StdD sequence from *S. thermodiastaticus* JCM 4840. Precursor sequences were identified through a manual search of the genome's region upstream of each StdE homolog.

| Gene ID | Strain | Phylum | %ID (MNIO/StdE) | Precursor peptide sequence |
|---|---|---|---|---|
| 8109937980 | *Streptomyces thermodiastaticus* JCM 4840 | Actinomycetota | 299/299 100% | MSDNTAAVLSAIDSDKAL RSAVIRHLATTHAHEFVG PLRAEFRGNDDIALASVN SNKEWQE |
| 2951696985 | *Actinomycetota* bacterium 052637 | Actinomycetota | 295/299 i98.66% | MSDNTAAVLSAIDSDKAL RSAVIRHLATTHAHEFVG PLRAEFRGNDDIALASVN SNKEWQE |
| 8112178169 | *Streptomyces thermoviolaceus apingens* JCM 4312 | Actinomycetota | 295/299 98.66% | MSDNTAAVLSAIDSDKAL RSAVIRHLATTHAHEFVG PLRAEFRGNDDIALASVN SNKEWQE |
| 8112201871 | *Streptomyces thermoviolaceus thermoviolaceus* JCM 4843 | Actinomycetota | 295/299 98.66% | MSDNTAAVLSAIDSDKAL RSAVIRHLATTHAHEFVG PLRAEFRGNDDIALASVN SNKEWQE |
| 2866522241 | *Streptomyces* sp. WAC00469 | Actinomycetota | 295/299 98.66% | MSDNTAAVLSAIDSDKAL RSAVIRHLATTHAHEFVG PLRAEFRGNDDIALASVN SNKEWQE |
| 2940111087 | *Actinomycetota* bacterium 52594 | Actinomycetota | 295/299 98.66% | MSDNTAAVLSAIDSDKAL RSAVIRHLATTHAHEFVG PLRAEFRGNDDIALASVN SNKEWQE |
| 2896974795 | *Streptomyces* sp. DSM 40868 | Actinomycetota | 278/299 92.98% | MSDNTAAVLSAIDSDKAL RSAVIRHLATTHANEFVS PLRAEFRGNDDIALASVN SNKEWQE |
| 2944895943 | *Streptomyces* sp. 25648 | Actinomycetota | 278/299 92.98% | MSDNTAAVLSAIDSDKAL RSAVIRHLATTHANEFVS PLRAEFRGNDDIALASVN SNKEWQE |
| 8012369728 | *Actinomycetota* bacterium 017043_259_D4 | Actinomycetota | 278/299 92.98% | MSDNTAAVLSAIDSDKAL RSAVIRHLATTHANEFVS PLRAEFRGNDDIALASVN SNKEWQE |
| 2975123176 | *Actinomycetota* bacterium 21046_187_C2 | Actinomycetota | 278/299 92.98% | MSDNTAAVLSAIDSDKAL RSAVIRHLATTHANEFVS PLRAEFRGNDDIALASVN SNKEWQE |
| 2960294871 | *Actinomycetota* bacterium 21112 | Actinomycetota | 278/299 92.98% | MSDNTAAVLSAIDSDKAL RSAVIRHLATTHANEFVS PLRAEFRGNDDIALASVN SNKEWQE |
| 2974034638 | *Actinomycetota* bacterium 22290 | Actinomycetota | 278/299 92.98% | MSDNTAAVLSAIDSDKAL RSAVIRHLATTHANEFVS PLRAEFRGNDDIALASVN SNKEWQE |
| 2766392705 | *Streptomyces lavenduligriseus* NRRL ISP-5487 | Actinomycetota | 278/299 92.98% | MSDNTAAVLSAIDSDKAL RSAVIRHLATTHANEFVS PLRAEFRGNDDIALASVN SNKEWQE |

| | | | | |
|---|---|---|---|---|
| 2945369264 | Actinomycetota bacterium 001328 | Actinomycetota | 278/299 92.98% | MSDKTAAVLSAIDSDKAL RSAVIRHLATTHANEFVS PLRAEFRGNDDIALASVN SNKEWQE |
| 8113491221 | *Streptomyces eurythermus* JCM 4206 | Actinomycetota | 275/299 91.97% | MSDNTAAVLSAIDSDKAL RSAVIRHLATTHANEFVS PLRAEFRGNDDIALASVN SNKEWQE |
| 8012981714 | *Actinomycetota* bacterium 014880_285_H06 | Actinomycetota | 275/299 91.97% | MSDNTAAVLSAIDSDKAL RSAVIRHLATTHANEFVS PLRAEFRGNDDIALASVN SNKEWQE |
| 8012350675 | *Actinomycetota* bacterium 014664_259_C8 | Actinomycetota | 275/299 91.97% | MSDNTAAVLSAIDSDKAL RSAVIRHLATTHANEFVS PLRAEFRGNDDIALASVN SNKEWQE |
| 8015150593 | *Actinomycetota* bacterium 014665_263_C06 | Actinomycetota | 275/299 91.97% | MSDNTAAVLSAIDSDKAL RSAVIRHLATTHANEFVS PLRAEFRGNDDIALASVN SNKEWQE |
| 8056831951 | *Streptomyces barringtoniae* JA03 | Actinomycetota | 238/299 79.60% | MSDKTTAILAALDGDKTL RSAIIQHLATTHAQEFVNP LRAELGGNDDLSLGAVNS NKEWQE |
| 2856761647 | *Streptacidiphilus pinicola* MMS16-CNU450 | Actinomycetota | 202/281 71.89% | MAEINEILAAIDSSKKVRS AVIERLVSTHAQEVVNPL RAIAGDDVSWSSITSNKD WSE |
| 2775746310 | *Myxococcales* bacterium CPC88 | Myxococcota | 131/276 47.46% | MVEALSKKLKGLSFEDLA EVIAEDREFREALIKHIAK DHAEDYLGEVNVNARVL AGLSARVVSSNKEWQE |
| 2775746310 | *Myxococcales* bacterium CPC88 | Myxococcota | 131/276 47.46% | MVLVGSKKLKGISFEEITA LIDEDREFREAMIRLLARD HAEDYLKQVNLNVGALTE LSARVVSSAKEWQD |
| 2775845471 | *Sandaracinus* sp. NAT131 | Myxococcota | 131/276 47.46% | MVEALSKKLKGLSFEDLA EVIAEDREFREALIKHIAK DHAEDYLGEVNVNARVL AGLSARVVSSNKEWQE |
| 2775845471 | *Sandaracinus* sp. NAT131 | Myxococcota | 131/276 47.46% | MVLVGSKKLKGISFEEITA LIDEDREFREAMIRLLARD HAEDYLKQVNLNVGALTE LSARVVSSAKEWQD |
| 2775851308 | *Sandaracinus* sp. SP2993 | Myxococcota | 131/276 47.46% | MVEALSKKLKGLSFEDLA EVIAEDREFREALIKHIAK DHAEDYLGEVNVNARVL AGLSARVVSSNKEWQE |
| 2775851308 | *Sandaracinus* sp. SP2993 | Myxococcota | 131/276 47.46% | MVLVGSKKLKGISFEEITA LIDEDREFREAMIRLLARD HAEDYLKQVNLNVGALTE LSARVVSSAKEWQD |
| 2982850127 | *Dickeya fangzhongdai* M074 | Pseudomonadota | 121/276 43.84% | MDKAQELELAVAELNAEN TEAENEILLDVTCMVSNK EWQE |
| 2982845965 | *Dickeya fangzhongdai* M005 | Pseudomonadota | 121/276 43.84% | MDKAQELELAVAELNAEN TEAENEILLDVTCMVSNK EWQE |
| 2630173480 | *Dickeya fangzhongdai* ND14b | Pseudomonadota | 121/276 43.84% | MDKAQELELAVAELNAEN TEAENEILLDVTCMVSNK EWQE |

| | | | | |
|---|---|---|---|---|
| 2776528989 | *Dickeya fangzhongdai* DSM 101947 | Pseudomonadota | 121/277<br>43.68% | MDKAQELELAVAELNAEN<br>TEAENEILLDVTCMVSNK<br>EWQE |
| 8113173844 | *Dickeya fangzhongdai* CGMCC 1.15464 | Pseudomonadota | 121/277<br>43.68% | MDKAQELELAVAELNAEN<br>TEAENEILLDVTCMVSNK<br>EWQE |
| 2860871481 | *Pseudomonas* sp. R1-43-08 | Pseudomonadota | 122/280<br>43.57% | MELQEKDVALEEQELAIA<br>KSEELSEELVLSLAHHAS<br>DKEWQE |
| 2651807005 | *Dickeya fangzhongdai* B16 | Pseudomonadota | 120/276<br>43.48% | MDKAQELELAVAELNAEN<br>TEAENEILLDVTCMVSNK<br>EWQE |
| 2559184145 | *Dickeya zeae* NCPPB 3531 | Pseudomonadota | 120/276<br>43.48% | MDKAQELELAVAELNAEN<br>TEAENEILLDVTCMVSNK<br>EWQE |
| 2654976169 | *Dickeya fangzhongdai* S1 | Pseudomonadota | 120/276<br>43.48% | MDKAQELELAVAELNAEN<br>TEAENEILLDVTCMVSNK<br>EWQE |
| 2814988251 | *Dickeya dadantii* Secpp1600 | Pseudomonadota | 120/276<br>43.48% | MDKAQELELAVAELNAEN<br>TEAENEILLDVTCMVSNK<br>EWQE |
| 2831161822 | *Dickeya zeae* A5272 | Pseudomonadota | 120/276<br>43.48% | MDKAQELELAVAELNAEN<br>TEAENEILLDVTCMVSNK<br>EWQE |
| 2982856433 | *Dickeya zeae* A5272 | Pseudomonadota | 120/276<br>43.48% | MDKAQELELAVAELNAEN<br>TEAENEILLDVTCMVSNK<br>EWQE |
| 2847241304 | *Dickeya fangzhongdai* PA1 | Pseudomonadota | 120/276<br>43.48% | MDKAQELELAVAELNAEN<br>TEAENEILLDVTCMVSNK<br>EWQE |
| 2559208148 | *Dickeya* sp. MK7 | Pseudomonadota | 120/276<br>43.48% | MDKAQELELAVAELNAEN<br>TEAENEILLDVTCMVSNK<br>EWQE |
| 2982835375 | *Dickeya zeae* A5410 | Pseudomonadota | 119/276<br>43.12% | MDKAQELELAVAELNAEN<br>AEAENEILLDVTCTVSNKE<br>WQE |
| 2559192607 | *Dickeya zeae* MK19 | Pseudomonadota | 119/276<br>43.12% | MDKAQELELAVAELNAEN<br>AEAENEILLDVTCTVSNKE<br>WQE |
| 2599197678 | *Dickeya zeae* NCPPB 3532 | Pseudomonadota | 119/276<br>43.12% | MDKAQELELAVAELNAEN<br>AEAENEILLDVTCTVSNKE<br>WQE |
| 2770339707 | *Dickeya zeae* AG740 | Pseudomonadota | 119/276<br>43.12% | MDKAQELELAVAELNAEN<br>AEAENEILLDVTCTVSNKE<br>WQE |
| 2843877218 | *Dickeya zeae* MS2 | Pseudomonadota | 119/276<br>43.12% | MDKAQELELAVAELNAEN<br>AEAENEILLDVTCTVSNKE<br>WQE |
| 2831156921 | *Dickeya zeae* A5410 | Pseudomonadota | 119/276<br>43.12% | MDKAQELELAVAELNAEN<br>AEAENEILLDVTCTVSNKE<br>WQE |
| 2559179939 | *Dickeya zeae* NCPPB 2538 | Pseudomonadota | 119/276<br>43.12% | MDKAQELELAVAELNAEN<br>AEAENEILLDVTCTVSNKE<br>WQE |
| 2982838804 | *Dickeya zeae* PL65 | Pseudomonadota | 119/276<br>43.12% | MDKAQELELAVAELNAEN<br>AEAENEILLDVTCTVSNKE<br>WQE |
| 646448342 | *Dickeya parazeae* Ech586 | Pseudomonadota | 118/276<br>42.75% | MDKAQELELAVAELNAEN<br>AEAENEILLDVTCTVSNKE<br>WQE |

| | | | | |
|---|---|---|---|---|
| 8085543270 | *Burkholderia* sp. MS455 | Pseudomonadota | 119/279 42.65% | MVEQNQFILEEDEKQIDANQVSDEELTLALSSSLSNKEWQE |
| 2552850907 | *Dickeya solani* MK10 | Pseudomonadota | 117/275 42.55% | MDKAQELELAVAELNAENTEAENEILLDVTCMVSNKEWQE |
| 2849505397 | *Photorhabdus* sp. HUG-39 | Pseudomonadota | 117/275 42.55% | MDKAQELELAVAELNAENADVENELLLDVASGVSNKEWQE |
| 2847232563 | *Dickeya solani* IFB0223 | Pseudomonadota | 117/275 42.55% | MDKAQELELAVAELNAENTEAENEILLDVTCMVSNKEWQE |
| 2854530711 | *Dickeya solani* IFB_0158 | Pseudomonadota | 117/275 42.55% | MDKAQELELAVAELNAENTEAENEILLDVTCMVSNKEWQE |
| 2895735474 | *Photorhabdus kayaii* C-HU2 | Pseudomonadota | 117/275 42.55% | MDKAQELELAVAELNAENADVENELLLDVASGVSNKEWQE |
| 2547655134 | *Dickeya solani* GBBC 2040 | Pseudomonadota | 117/275 42.55% | MDKAQELELAVAELNAENTEAENEILLDVTCMVSNKEWQE |
| 2843868778 | *Dickeya solani* D s0432-1 | Pseudomonadota | 117/275 42.55% | MDKAQELELAVAELNAENTEAENEILLDVTCMVSNKEWQE |
| 2847245787 | *Dickeya solani* IFB 0099 | Pseudomonadota | 117/275 42.55% | MDKAQELELAVAELNAENTEAENEILLDVTCMVSNKEWQE |
| 2841397997 | *Dickeya solani* RNS 08.23.3.1.A | Pseudomonadota | 117/275 42.55% | MDKAQELELAVAELNAENTEAENEILLDVTCMVSNKEWQE |
| 2565704031 | *Dickeya solani* D s0432-1 | Pseudomonadota | 117/275 42.55% | MDKAQELELAVAELNAENTEAENEILLDVTCMVSNKEWQE |
| 2552837950 | *Dickeya solani* MK16 | Pseudomonadota | 117/275 42.55% | MDKAQELELAVAELNAENTEAENEILLDVTCMVSNKEWQE |
| 2895749269 | *Photorhabdus kayaii* M-HU2 | Pseudomonadota | 117/275 42.55% | MDKAQELELAVAELNAENADVENELLLDVASGVSNKEWQE |
| 2684803920 | *Dickeya solani* RNS 07.7.3B | Pseudomonadota | 117/275 42.55% | MDKAQELELAVAELNAENTEAENEILLDVTCMVSNKEWQE |
| 2854598718 | *Dickeya solani* PPO 9019 | Pseudomonadota | 117/275 42.55% | MDKAQELELAVAELNAENTEAENEILLDVTCMVSNKEWQE |
| 2673345593 | *Dickeya solani* PPO 9134 | Pseudomonadota | 117/275 42.55% | MDKAQELELAVAELNAENTEAENEILLDVTCMVSNKEWQE |
| 2841387782 | *Dickeya solani* IPO 2222 | Pseudomonadota | 117/275 42.55% | MDKAQELELAVAELNAENTEAENEILLDVTCMVSNKEWQE |
| 2854615059 | *Dickeya solani* IFB_0221 | Pseudomonadota | 117/275 42.55% | MDKAQELELAVAELNAENTEAENEILLDVTCMVSNKEWQE |
| 2552833607 | *Dickeya* solani IPO 2222 | Pseudomonadota | 117/275 42.55% | MDKAQELELAVAELNAENTEAENEILLDVTCMVSNKEWQE |
| 2834863738 | *Dickeya dianthicola* WV516 | Pseudomonadota | 117/276 42.39% | MDKAQELELAVAELNAENTESENEILLDVACTVSNKEWQE |

| | | | | |
|---|---|---|---|---|
| 2552855238 | *Dickeya dianthicola* IPO 980 | Pseudomonadota | 117/276 42.39% | MDKAQELELAVAELNAEN TESENEILLDVACTVSNKE WQE |
| 2847228062 | *Dickeya dianthicola* ME23 | Pseudomonadota | 117/276 42.39% | MDKAQELELAVAELNAEN TESENEILLDVACTVSNKE WQE |
| 2854576163 | *Dickeya dianthicola* DE440 | Pseudomonadota | 117/276 42.39% | MDKAQELELAVAELNAEN TESENEILLDVACTVSNKE WQE |
| 8079946014 | *Dickeya dianthicola* LAR.16.03.LID | Pseudomonadota | 117/276 42.39% | MDKAQELELAVAELNAEN TESENEILLDVACTVSNKE WQE |
| 2552859749 | *Dickeya dianthicola* NCPPB 3534 | Pseudomonadota | 117/276 42.39% | MDKAQELELAVAELNAEN TESENEILLDVACTVSNKE WQE |
| 2630773784 | *Dickeya dianthicola* RNS04.9 | Pseudomonadota | 117/276 42.39% | MDKAQELELAVAELNAEN TESENEILLDVACTVSNKE WQE |
| 2552841937 | *Dickeya dianthicola* GBBC 2039 | Pseudomonadota | 117/276 42.39% | MDKAQELELAVAELNAEN TESENEILLDVACTVSNKE WQE |
| 2552846655 | *Dickeya dianthicola* NCPPB 453 | Pseudomonadota | 117/276 42.39% | MDKAQELELAVAELNAEN TESENEILLDVACTVSNKE WQE |
| 2854609977 | *Dickeya dianthicola* S4.16.03.P2.4 | Pseudomonadota | 117/276 42.39% | MDKAQELELAVAELNAEN TESENEILLDVACTVSNKE WQE |
| 2559220795 | *Dickeya* sp. CSL RW240 | Pseudomonadota | 117/276 42.39% | MDKAQELELAVAELNAEN TESENEILLDVACTVSNKE WQE |
| 2854602561 | *Dickeya dianthicola* SS70 | Pseudomonadota | 117/276 42.39% | MDKAQELELAVAELNAEN TESENEILLDVACTVSNKE WQE |
| 2854587708 | *Dickeya dianthicola* S4.16.03.LID | Pseudomonadota | 117/276 42.39% | MDKAQELELAVAELNAEN TESENEILLDVACTVSNKE WQE |
| 2981433753 | *Paraburkholderia* sp. GAS199 | Pseudomonadota | 117/277 42.24% | MNEEMQSALEQDAKVLE AEQIAEDELTLSLSQLISN KEWQE |
| 8002953181 | *Dickeya solani* RNS 05.1.2A | Pseudomonadota | 116/275 42.18% | MDKAQKLELAVAELNAEN TEAENEILLDVTCMVSNK EWQE |
| 2559216572 | *Dickeya* sp. NCPPB 3274 | Pseudomonadota | 116/275 42.18% | MDKAQELELAVAELNAEN TEAENEILLDVTCMVSNK EWQE |
| 8044042700 | *Pseudomonas coronafaciens* pv. *atropurpurea* ICMP 4451 | Pseudomonadota | 118/280 42.14% | MELQEKDVALEEQELAIA KSEELSEELVLSLAHHGS DKEWQE |
| 2834855307 | *Dickeya undicola* FVG10-MFV-A16 | Pseudomonadota | 115/276 41.67% | MDKAQELELAVAELNAEN AEAENEILLDVTCMVSNK EWQE |
| 8064555015 | *Yokenella regensburgei* NCTC 12131 | Pseudomonadota | 116/280 41.43% | MDKSQELELAVAELNAEN AEAENELLLDVTCLVSNK EWQE |
| 2757616447 | *Yokenella regensburgei* DSM 5079 | Pseudomonadota | 116/280 41.43% | MDKSQELELAVAELNAEN AEAENELLLDVTCLVSNK EWQE |
| 8102243379 | *Caballeronia* sp. LZ043 | Pseudomonadota | 115/278 41.37% | MIMNEQNQAVLENDVRQI DVEQVADDELTLALSHVV SNKEWQE |

| | | | | |
|---|---|---|---|---|
| 8025682312 | *Caballeronia hypogeia* LZ043 | Pseudomonadota | 115/278 41.37% | MIMNEQNQAVLENDVRQIDVEQVADDELTLALSHVVSNKEWQE |
| 2629162083 | *Dickeya undicola* 2B12 | Pseudomonadota | 114/276 41.30% | MDKAQELELAVAELNAENAEAENEILLDVTCMVSNKEWQE |
| 2773180829 | *Xenorhabdus ehlersii* DSM 16337 | Pseudomonadota | 113/275 41.09% | MDKAQELEIAVAELNAENSDAENELLLDITGVVSNKEWQE |
| 2839587921 | *Xenorhabdus ishibashii* DSM 22670 | Pseudomonadota | 113/275 41.09% | MDKAQELEIAVAELNAENSDAENELLLDITGVVSNKEWQE |
| 8025655193 | *Caballeronia hypogeia* LZ032 | Pseudomonadota | 114/278 41.01% | MIMNEQNQAVLENDVRQIDVEQVTDDELTLALSHVVSNKEWQE |
| 8102212807 | *Caballeronia* sp. LZ032 | Pseudomonadota | 114/278 41.01% | MIMNEQNQAVLENDVRQIDVEQVTDDELTLALSHVVSNKEWQE |
| 2769553685 | *Pseudomonas* sp. 37 R 15 | Pseudomonadota | 103/276 37.32% | MDSQQATQLELDEQLVEVDLHAEVDDELVFAISAPTSNKEWQE |
| 2769519119 | *Pseudomonas* sp. 28 E 9 | Pseudomonadota | 103/276 37.32% | MDSQQATQLELDEQLVEVDLHAEVDDELVLAISAPTSNKEWQE |

**Table S3.** HR-MS/MS fragmentation data for Arg-C Ultra fragment of unmodified StdA.

| Ion | Observed *m/z* | Calculated *m/z* | Δ ppm | Sequence |
|---|---|---|---|---|
| $b_3$ | 287.0988 | 287.0986 | 0.0002 | GND |
| $b_4$ | 402.1255 | 402.1256 | 0.0001 | GNDD |
| $b_5$ | 515.2092 | 515.2096 | 0.0004 | GNDDI |
| $b_6$ | 586.2464 | 586.2467 | 0.0003 | GNDDIA |
| $b_7$ | 699.3309 | 699.3308 | 0.0001 | GNDDIAL |
| $b_8$ | 770.3674 | 770.3679 | 0.0005 | GNDDIALA |
| $b_9$ | 857.3995 | 857.3999 | 0.0004 | GNDDIALAS |
| $b_{10}$ | 956.4703 | 956.4683 | 0.002 | GNDDIALASV |
| $b_{11}$ | 1070.5123 | 1070.5113 | 0.001 | GNDDIALASVN |
| $b_{12}$ | 1157.5332 | 1157.5433 | 0.0101 | GNDDIALASVNS |
| $b_{13}$ | 1271.5896 | 1271.5862 | 0.0034 | GNDDIALASVNSN |
| $b_{14}$ | 1399.6791 | 1399.6812 | 0.0021 | GNDDIALASVNSNK |
| $b_{15}$ | 1528.7231 | 1528.7238 | 0.0007 | GNDDIALASVNSNKE |
| $b_{16}$ | 1714.7986 | 1714.8031 | 0.0045 | GNDDIALASVNSNKEW |
| $y_2$ | 276.1191 | 276.1190 | 0.0001 | QE |
| $y_3$ | 462.1979 | 462.1983 | 0.0004 | WQE |
| $y_4$ | 591.2402 | 591.2409 | 0.0007 | EWQE |
| $y_5$ | 719.3354 | 719.3359 | 0.0005 | KEWQE |
| $y_6$ | 833.3771 | 833.3788 | 0.0017 | NKEWQE |
| $y_7$ | 920.4091 | 920.4108 | 0.0017 | SNKEWQE |
| $y_8$ | 1034.4514 | 1034.4538 | 0.0024 | NSNKEWQE |
| $y_9$ | 1133.5203 | 1133.5222 | 0.0019 | VNSNKEWQE |
| $y_{10}$ | 1220.5522 | 1220.5542 | 0.002 | SVNSNKEWQE |
| $y_{11}$ | 1291.5896 | 1291.5913 | 0.0017 | ASVNSNKEWQE |
| $y_{12}$ | 1404.6770 | 1404.6754 | 0.0016 | LASVNSNKEWQE |
| $y_{13}$ | 1475.7136 | 1475.7125 | 0.0011 | ALASVNSNKEWQE |
| $y_{14}$ | 1588.7948 | 1588.7966 | 0.0018 | IALASVNSNKEWQE |
| $y_{15}$ | 1703.8313 | 1703.8235 | 0.0078 | DIALASVNSNKEWQE |
| $y_{16}$ | 1818.8472 | 1818.8504 | 0.0032 | DDIALASVNSNKEWQE |

**Table S4.** HR-MS/MS data for Arg-C Ultra fragment of StdA following the in vitro reaction with StdGH in the presence of ATP and $Na_2S$.

| Ion | Observed m/z | Calculated m/z | Δ ppm | Sequence |
|---|---|---|---|---|
| $b_3$ | 287.0990 | 287.0986 | 0.0004 | GND |
| $b_4$ | 402.1256 | 402.1256 | 0 | GNDD |
| $b_5$ | 515.2098 | 515.2096 | 0.0002 | GNDDI |
| $b_6$ | 586.2473 | 586.2467 | 0.0006 | GNDDIA |
| $b_7$ | 699.3293 | 699.3308 | 0.0015 | GNDDIAL |
| $b_8$ | 770.3698 | 770.3679 | 0.0019 | GNDDIALA |
| $b_9$ | 857.4064 | 857.3999 | 0.0065 | GNDDIALAS |
| $b_{10}$ | 956.4700 | 956.4683 | 0.0017 | GNDDIALASV |
| $b_{11}$ | 1070.5110 | 1070.5113 | 0.0003 | GNDDIALASVN |
| $b_{12}$ | 1157.5421 | 1157.5433 | 0.0012 | GNDDIALASVNS |
| $b_{14}$ | 1415.6562 | 1415.6512 | 0.005 | GNDDIALASVNSnK |
| $b_{15}$ | 1544.7006 | 1544.6938 | 0.0068 | GNDDIALASVNSnKE |
| $b_{16}$ | 1730.782 | 1730.7731 | 0.0089 | GNDDIALASVNSnKEW |
| $y_2$ | 276.1193 | 276.1190 | 0.0003 | QE |
| $y_3$ | 462.1983 | 462.1983 | 0 | WQE |
| $y_4$ | 591.2411 | 591.2409 | 0.0002 | EWQE |
| $y_5$ | 719.3402 | 719.3359 | 0.0043 | KEWQE |
| $y_6$ | 849.3528 | 849.3488 | 0.004 | nKEWQE |
| $y_7$ | 936.3867 | 936.3808 | 0.0059 | SnKEWQE |
| $y_8$ | 1050.4291 | 1050.4238 | 0.0053 | NSnKEWQE |
| $y_9$ | 1149.4984 | 1149.4922 | 0.0062 | VNSnKEWQE |
| $y_{10}$ | 1236.5306 | 1236.5242 | 0.0064 | SVNSnKEWQE |
| $y_{11}$ | 1307.5674 | 1307.5613 | 0.0061 | ASVNSnKEWQE |
| $y_{12}$ | 1420.6522 | 1420.6454 | 0.0068 | LASVNSnKEWQE |
| $y_{13}$ | 1491.6895 | 1491.6825 | 0.007 | ALASVNSnKEWQE |
| $y_{14}$ | 1604.7765 | 1604.7666 | 0.0099 | IALASVNSnKEWQE |

**Table S5.** HR-MS/MS data for Arg-C Ultra fragment of StdGH-modified StdA peptide following the in vitro reaction with StdF in the presence of carbamoyl phosphate and ATP. This species represents the carbamoylated full-length peptide.

| Ion | Observed m/z | Calculated m/z | Δ ppm | Sequence |
|---|---|---|---|---|
| $b_3$ | 287.0988 | 287.0986 | 0.0002 | GND |
| $b_4$ | 402.1251 | 402.1256 | 0.0005 | GNDD |
| $b_5$ | 515.2084 | 515.2096 | 0.0012 | GNDDI |
| $b_6$ | 586.2462 | 586.2467 | 0.0005 | GNDDIA |
| $b_7$ | 699.3296 | 699.3308 | 0.0012 | GNDDIAL |
| $b_8$ | 770.3666 | 770.3679 | 0.0013 | GNDDIALA |
| $b_9$ | 857.3995 | 857.3999 | 0.0004 | GNDDIALAS |
| $b_{10}$ | 956.4658 | 956.4683 | 0.0025 | GNDDIALASV |
| $b_{11}$ | 1070.5096 | 1070.5113 | 0.0017 | GNDDIALASVN |
| $b_{12}$ | 1157.5464 | 1157.5433 | 0.0031 | GNDDIALASVNS |
| $b_{13}$ | 1287.5669 | 1287.5562 | 0.0107 | GNDDIALASVNSn |
| $b_{14}$ | 1458.6632 | 1458.6570 | 0.0062 | GNDDIALASVNSnk |
| $y_2$ | 276.119 | 276.1190 | 0 | QE |
| $y_3$ | 462.1977 | 462.1983 | 0.0006 | WQE |
| $y_4$ | 591.2402 | 591.2409 | 0.0007 | EWQE |
| $y_5$ | 762.3392 | 762.3417 | 0.0025 | kEWQE |
| $y_8$ | 1093.4354 | 1093.4296 | 0.0058 | NSnkEWQE |

**Table S6.** HR-MS/MS data for Arg-C Ultra fragment for cleaved StdGH-modified StdA peptide following the in vitro reaction with StdF in the presence of carbamoyl phosphate and ATP, and StdDE in the presence of iron and ascorbate.

| Ion | Observed *m/z* | Calculated *m/z* | Δ ppm | Sequence |
|---|---|---|---|---|
| $b_3$ | 287.0995 | 287.0986 | 0.0009 | GND |
| $b_4$ | 402.1265 | 402.1256 | 0.0009 | GNDD |
| $b_5$ | 515.2108 | 515.2096 | 0.0012 | GNDDI |
| $b_6$ | 586.2482 | 586.2467 | 0.0015 | GNDDIA |
| $b_7$ | 699.3339 | 699.3308 | 0.0031 | GNDDIAL |
| $b_8$ | 770.3704 | 770.3679 | 0.0025 | GNDDIALA |
| $b_9$ | 857.4037 | 857.3999 | 0.0038 | GNDDIALAS |
| $b_{12}$ | 1157.5503 | 1157.5433 | 0.007 | GNDDIALASVNS |
| $y_1$ | 222.1092 | 222.1084 | 0.0008 | k |
| $y_2$ | 352.1296 | 352.1213 | 0.0083 | nk |
| $y_3$ | 439.1618 | 439.1534 | 0.0084 | Snk |
| $y_4$ | 553.2064 | 553.1963 | 0.0101 | NSnk |
| $y_6$ | 739.3060 | 739.2967 | 0.0093 | SVNSnk |
| $y_7$ | 810.3428 | 810.3338 | 0.009 | ASVNSnk |
| $y_8$ | 923.4227 | 923.4179 | 0.0048 | LASVNSnk |
| $y_9$ | 994.4649 | 994.4550 | 0.0099 | ALASVNSnk |

**Table S7.** Plasmids and bacterial strains used in this study.

| Plasmids | Purpose | Source |
|---|---|---|
| pRSFDuet-1_*stdH_stdG* | $Kan^R$, Coexpression of StdG and StdH | This study |
| pRSFDuet-1_*stdF* | $Kan^R$, Coexpression of StdG and StdGH | This study |
| pRSFDuet-1_*stdD_stdE* | $Kan^R$, Coexpression of StdD and StdE | This study |
| **Bacterial strains** | **Purpose** | **Source** |
| *E. coli* BL21 (DE3) | Coexpression of StdDE and expression of StdF | NEB |
| *E. coli* Rosetta2 (DE3) pLysS | $CM^R$, Coexpression of StdGH | NEB |

**Table S8.** LC-MS ESI data for StdA following the in vitro reaction with StdGH in the presence of ATP and $Na_2S$.

| Substrate | Enzyme | Product | Ion | Observed *m/z* | Calculated *m/z* | Δ ppm |
|---|---|---|---|---|---|---|
| StdA | No enzyme | StdA | $[M+10H]^{10+}$ | 665.8316 | 665.7290 | 0.1026 |
| | | | $[M+9H]^{9+}$ | 739.7006 | 739.5878 | 0.1128 |
| | | | $[M+8H]^{8+}$ | 832.0387 | 831.9113 | 0.1274 |
| | | | $[M+7H]^{7+}$ | 950.7580 | 950.6129 | 0.1451 |
| | | | $[M+6H]^{6+}$ | 1109.0518 | 1108.8817 | 0.1701 |
| | | | $[M+5H]^{5+}$ | 1330.6643 | 1330.4480 | 0.2163 |
| | | | $[M+4H]^{4+}$ | 1663.0825 | 1662.8225 | 0.26 |
| StdA | StdGH | StdGH-modified StdA | $[M+10H]^{10+}$ | 667.4294 | 667.3240 | 0.1054 |
| | | | $[M+9H]^{9+}$ | 741.4731 | 741.3600 | 0.1131 |
| | | | $[M+8H]^{8+}$ | 834.0325 | 833.9050 | 0.1275 |
| | | | $[M+7H]^{7+}$ | 953.0361 | 952.8914 | 0.1447 |
| | | | $[M+6H]^{6+}$ | 1111.7092 | 1111.5400 | 0.1692 |
| | | | $[M+5H]^{5+}$ | 1333.8506 | 1333.6480 | 0.2026 |

**Table S9.** LC-MS ESI Data for StdGH-modified StdA following in vitro reactions with (1) StdDE in the presence of iron and ascorbate and (2) StdF in the presence of carbamoyl phosphate and ATP.

| **Substrate** | **Enzyme** | **Product** | **Ion** | **Observed *m/z*** | **Calculated *m/z*** | **Δ ppm** |
|---|---|---|---|---|---|---|
| StdGH-modified StdA | StdDE | StdGH-modified StdA | $[M+10H]^{10+}$ | 667.4294 | 667.3240 | 0.1054 |
| | | | $[M+9H]^{9+}$ | 741.4731 | 741.3600 | 0.1131 |
| | | | $[M+8H]^{8+}$ | 834.0325 | 833.9050 | 0.1275 |
| | | | $[M+7H]^{7+}$ | 953.0361 | 952.8914 | 0.1447 |
| | | | $[M+6H]^{6+}$ | 1111.7092 | 1111.5400 | 0.1692 |
| | | | $[M+5H]^{5+}$ | 1333.8506 | 1333.6480 | 0.2026 |
| StdGH-modified StdA | StdF | Full-length StdGHF-modified StdA | $[M+9H]^{9+}$ | 746.1476 | 746.1311 | 0.0165 |
| | | | $[M+8H]^{8+}$ | 839.2907 | 839.2725 | 0.0182 |
| | | | $[M+7H]^{7+}$ | 959.0450 | 959.0257 | 0.0193 |
| | | | $[M+6H]^{6+}$ | 1118.7221 | 1118.6967 | 0.0254 |
| | | | $[M+5H]^{5+}$ | 1342.2653 | 1342.2360 | 0.0293 |
| StdGH-modified StdA | StdF | Cleaved StdGHF-modified StdA | $[M+9H]^{9+}$ | 682.5701 | 682.5533 | 0.0168 |
| | | | $[M+8H]^{8+}$ | 767.7634 | 767.7475 | 0.0159 |
| | | | $[M+7H]^{7+}$ | 877.3003 | 877.2829 | 0.0174 |
| | | | $[M+6H]^{6+}$ | 1023.3502 | 1023.3300 | 0.0202 |
| | | | $[M+5H]^{5+}$ | 1227.8215 | 1227.7960 | 0.0255 |

**Table S10.** LC-MS ESI Data for StdGHF-modified StdA following in vitro reactions with StdDE in the presence of iron and ascorbate.

| Substrate | Enzyme | Product | Ion | Observed m/z | Calculated m/z | Δ ppm |
|---|---|---|---|---|---|---|
| Cleaved StdGHF-modified StdA | StdDE | Cleaved StdGHFDE-modified StdA (mono-hydroxylation) | $[M+9H]^{9+}$ | 684.3317 | 684.3433 | 0.0116 |
| | | | $[M+8H]^{8+}$ | 769.7441 | 769.7612 | 0.0171 |
| | | | $[M+7H]^{7+}$ | 879.5641 | 879.5943 | 0.0302 |
| | | | $[M+6H]^{6+}$ | 1025.9902 | 1026.0150 | 0.0248 |
| | | | $[M+5H]^{5+}$ | 1230.9912 | 1231.0180 | 0.0268 |
| | | | $[M+4H]^{4+}$ | 1538.4871 | 1538.5225 | 0.0354 |
| Cleaved StdGHF-modified StdA | StdDE | Cleaved StdGHFDE-modified StdA (bis-hydroxylation) | $[M+8H]^{8+}$ | 771.8662 | 771.7612 | 0.105 |
| | | | $[M+7H]^{7+}$ | 881.9900 | 881.8700 | 0.12 |
| | | | $[M+6H]^{6+}$ | 1028.8221 | 1028.6817 | 0.1404 |
| | | | $[M+5H]^{5+}$ | 1234.3870 | 1234.2180 | 0.169 |
| | | | $[M+4H]^{4+}$ | 1542.7346 | 1542.5225 | 0.2121 |
| | | | $[M+3H]^{3+}$ | 2056.6452 | 2056.3633 | 0.2819 |

**Table S11.** HPLC retention times of Arg-C Ultra digested modified StdA products.

| ArgC-Ultra cleaved product | Retention time (min) |
|---|---|
| Unmodified StdA | 26.45 |
| StdGH-modified StdA | 17.65 |
| StdGHF-modified StdA (full-length) | 30.28 |
| StdGHFDE-modified StdA | 25.15 |

**DNA and Amino Acid Sequences**

**<u>StdA</u>**

*Amino acid sequence*

MSDNTAAVLSAIDSDKALRSAVIRHLATTHAHEFVGPLRAEFRGNDDIALASVNSNKEW
QE

*DNA sequence*

ATGTCCGACAACACCGCTGCGGTACTGTCCGCCATCGACAGCGACAAGGCGCTG
CGTTCGGCGGTCATCCGGCACCTCGCCACGACCCACGCCCACGAGTTCGTCGGT
CCGCTGCGCGCCGAGTTCCGCGGCAACGACGACATCGCCCTGGCGTCGGTCAAC
TCCAACAAGGAGTGGCAGGAGTAA

**<u>StdD (Multiple Cloning Site 1)</u>**

*Amino acid sequence*

MRTLSSGVTQALLLRDDREARTELLLRLLYSKSSADVVADALKDPSGRMCEATASVLS
TIDFHALEVERSARLRNLNRHLQRTCPPLWDCVEAETITPLVEAYADSDAFWQSAGRT
LFENFCLFAHRTVAGQSQLLADVFQLFGLVSRFSAPELIAPPEDVCACTTGIPGEGMDL
AKAAFSPWDDGGLIPPHVPDALQTESFRSAWRLVDERGRLPRSLDPDALGDPGDYQI
VVAAFPGRKVSAAALPLSGRCA

*DNA sequence*

TTGCGCACCCTTTCTTCCGGCGTGACGCAGGCACTCCTGTTGCGCGACGACAGAG
AGGCCCGTACGGAGTTGCTGCTCCGCCTGCTGTACTCCAAGAGCTCCGCGGACGT
GGTCGCCGATGCGCTGAAAGACCCCTCCGGCAGGATGTGCGAAGCAACGGCGTC
CGTTCTTTCCACCATCGATTTCCACGCCCTGGAGGTCGAGCGAAGCGCCCGTCTG
CGCAATCTGAACCGTCATCTGCAACGCACGTGCCCGCCTCTGTGGGATTGCGTCG
AAGCGGAAACGATCACTCCGCTCGTCGAGGCCTACGCGGACTCCGATGCTTTCTG
GCAGAGCGCCGGTCGCACCCTGTTCGAGAATTTCTGTCTGTTCGCCCACAGGACC
GTCGCCGGGCAGTCCCAGCTGCTCGCGGACGTCTTTCAGCTGTTCGGCCTCGTGA
GCCGCTTCTCCGCTCCCGAGTTGATCGCGCCTCCCGAGGACGTGTGTGCCTGCAC
GACCGGTATTCCGGGAGAGGGCATGGATCTGGCGAAGGCCGCCTTCTCCCCCTG
GGACGACGGCGGACTGATCCCCCCGCACGTTCCGGACGCCCTCCAGACCGAGTC
GTTCCGCTCTGCCTGGCGCCTCGTGGACGAGCGCGGGCGGCTGCCGCGCTCCCT
CGACCCCGACGCCCTCGGTGACCCCGGTGACTACCAGATCGTTGTCGCCGCCTTC
CCCGGCCGCAAGGTGTCCGCCGCGGCCCTTCCGCTCTCAGGCAGGTGCGCATGA

*DNA sequence (E. coli optimized)*

ATGAGGACATTATCAAGTGGAGTAACTCAAGCGTTGCTCCTACGCGATGACCGTGA
AGCTAGAACTGAGTTGCTGTTACGCTTGCTTTACAGCAAGTCGAGCGCCGATGTCG
TCGCCGACGCGCTGAAGGATCCGTCTGGTCGTATGTGCGAAGCTACGGCAAGCG
TTCTGAGTACCATCGATTTTCACGCTTTGGAAGTTGAACGTAGCGCGCGTTTGCGC
AACCTGAATCGTCATCTGCAACGTACCTGCCCGCCGCTGTGGGACTGTGTTGAGG
CCGAAACGATTACCCCGTTAGTAGAGGCGTATGCAGACAGCGATGCGTTCTGGCA
GTCGGCTGGCCGCACCCTGTTCGAGAACTTCTGCCTGTTCGCACACCGTACGGTT
GCTGGCCAGTCCCAGCTGTTGGCCGACGTGTTTCAACTGTTCGGCCTGGTGAGCC

GTTTTTCCGCGCCTGAGTTGATCGCTCCACCGGAGGATGTGTGCGCGTGCACCAC
CGGTATTCCGGGTGAAGGCATGGATCTGGCTAAAGCAGCGTTCTCTCCGTGGGAC
GACGGCGGTCTGATCCCGCCGCATGTTCCGGATGCACTGCAAACCGAGAGCTTTC
GTAGCGCGTGGCGTCTGGTTGATGAACGCGGTCGTCTCCCGCGTTCTCTGGATCC
AGACGCGCTGGGTGACCCGGGCGACTACCAGATTGTGGTGGCCGCGTTTCCGGG
TCGCAAAGTGTCAGCGGCAGCGCTGCCGCTGTCCGGTCGCTGTGCGTAA

**StdE (Multiple Cloning Site 2)**

*Amino acid sequence*

MTKEKKDPVGLLGFGLDWQYRDSYPQKMPDLCARYAGRLSHLSCVSLPSAADAQTF
VDECAGDLPVVHHLPGVAPAAPGGPNMELFTRLEAVSDVLGAVWTCEDIGLWSIGPY
PLPYFTPPLFEREIADHVVDGIRRMQEVSRYPFVPEIPSCTLAVGRMSLGEFFHRVTDG
ADCDLLLDVAHVFSYAVAVRQPYEDVLRSLPLDRVVEIHVAGGYIDPTFDNRYLDTHSH
AVTPAVIDLLQEAVAHAPRLRAVTYEIGVGLGADELDSDFERIENLLAEVGWTPSISRPS
GLATAGTA

*DNA sequence*

ATGACGAAAGAGAAGAAGGACCCCGTCGGCCTGCTCGGCTTCGGACTCGACTGG
CAGTACCGCGACAGCTACCCGCAGAAGATGCCGGACCTGTGCGCCCGGTACGCC
GGCCGGCTCAGCCATCTGTCCTGTGTCTCCCTGCCCAGCGCCGCCGACGCGCAG
ACCTTCGTGGACGAGTGCGCCGGCGACCTGCCTGTGGTGCACCACCTGCCGGGC
GTGGCACCCGCAGCGCCCGGCGGCCCCAACATGGAGCTGTTCACCCGCCTCGAG
GCGGTCAGCGACGTGCTGGGTGCCGTGTGGACCTGTGAGGACATCGGCCTGTGG
TCGATCGGCCCGTACCCGCTGCCGTACTTCACCCCTCCGCTCTTCGAGCGCGAGA
TCGCCGACCATGTCGTCGACGGCATACGCCGCATGCAGGAGGTCAGCCGTTACC
CCTTCGTGCCGGAGATCCCGTCCTGCACCCTCGCCGTGGGCCGTATGAGCCTCG
GCGAGTTCTTCCATCGCGTCACCGACGGCGCCGACTGCGACCTGCTGCTGGACG
TCGCGCATGTCTTCTCGTACGCCGTCGCCGTGCGGCAGCCCTACGAGGACGTGCT
CCGCTCGCTTCCGCTGGACCGGGTGGTGGAGATCCATGTGGCGGGCGGTTACAT
CGACCCGACCTTCGACAACCGGTACCTGGACACCCACAGCCACGCCGTGACCCC
GGCCGTCATCGACCTGCTCCAGGAGGCGGTCGCCCACGCGCCGCGGCTGCGCG
CCGTCACCTACGAGATCGGGGTGGGTCTCGGGGCCGACGAGCTGGACAGCGACT
TCGAGCGCATCGAGAACCTGCTGGCGGAGGTCGGGTGGACCCCGTCCATCTCCC
GTCCGTCCGGCCTGGCCACGGCAGGTACCGCATGA

*DNA sequence (E. coli optimized)*

ATGACAAAAGAAAAGAAAGATCCCGTAGGACTCTTGGGCTTCGGCCTCGACTGGC
AGTATCGCGATAGCTACCCGCAGAAAATGCCGGATCTTTGCGCGCGTTACGCTGG
CCGTCTGAGCCACCTGTCATGCGTTAGCCTGCCGTCTGCGGCGGATGCTCAAACC
TTTGTTGACGAGTGCGCAGGCGATCTGCCGGTTGTTCACCACCTGCCGGGTGTGG
CGCCAGCAGCGCCTGGTGGTCCGAATATGGAACTGTTCACGCGTCTGGAAGCGG
TCTCCGATGTTCTGGGCGCGGTTTGGACGTGCGAAGACATCGGCCTGTGGAGCAT
CGGCCCGTACCCGTTGCCGTACTTCACCCCACCGTTGTTTGAACGTGAAATCGCT
GACCACGTGGTGGATGGTATTCGCCGTATGCAAGAGGTGAGCCGTTACCCGTTTG
TCCCGGAGATCCCGTCTTGTACCCTGGCGGTTGGGCGTATGAGCCTGGGCGAATT

CTTTCATCGTGTTACCGACGGTGCTGACTGTGATTTGCTGCTGGACGTTGCGCACG
TGTTTAGCTATGCGGTGGCCGTACGCCAGCCGTACGAGGATGTTCTTAGATCGTT
GCCGTTGGACCGCGTAGTGGAAATTCATGTGGCGGGTGGCTATATTGATCCGACC
TTCGATAACCGTTATCTGGACACCCATTCCCACGCCGTGACTCCGGCAGTGATTGA
TTTACTGCAAGAGGCGGTCGCCCATGCACCGCGTCTGCGCGCAGTCACCTATGAG
ATCGGTGTTGGTCTGGGTGCTGACGAGCTGGACTCCGACTTCGAGCGCATTGAAA
ACCTGTTGGCCGAAGTTGGTTGGACTCCGTCGATCAGCCGTCCCAGCGGTCTGGC
TACCGCGGGCACGGCATAA

**StdF** **<u>(Multiple Cloning Site 1)</u>**

*Amino acid sequence*

MDPVHLPSVRPGHGRYRMTHAQQPASPPWSVLSPEPDRGAGLGEGVRVLVVSDGT
PAPDGDGMPDAHTERVAGLCRQLAGRAEIGYASAARLRARSLAADLRAMVTADGGW
DIVAMPWSSPERDAERLKVRRAFEEVARDPDMPLFVAAAGHDGPGRLRFPASCPSVL
AVGVGSDHGGPTAYCGTDGLGRKPQLLVPDARYATRCAAGPAPMRGTSAAVGIVAG
LAAVLAQRLGSDGRRVPSPLLRAALLASAGPDGTLVGTPLLRQPGTGAAEPFCFELPA
AAARLRLRLRTGADTVRIAAVAAGADEVVSGDTAPLWLPRAAELSVTWAADGTTRRTT
GAGWLVLDVASGAGRDTVITLECPEPVTAYVAVGGAEAVERIDPRLEKTDEPGVTAPA
WRAGTGRPVVLGLSASHDASACLLRDGSPQVAVQQERVTRRKHDGVGHLSSR
AAADYCLTSAGLTADDVDVFAFNAQPLLPGYVGLSVPSAAQSFDLFDPFDARTVYVSH
HLAHAFSAFWGSPFEEAVVVVCDGSGGSVLGTDDLLVDGPGLREYLARGPEDRLPRI
HVFSVYHFDRDGYRLLFRECADSFNVRVGSSSIGETYAAVSQYVFGDWQEGSGKLM
GLAPWGKARNAGPSLLEPGPHGLPRFRSDWKDTYRAPGSGGPMDHADLAARIQADL
EEALVARMRYAMSLVPGCRNLAYAGGVALNSVANDRIARESGADRFFVFPAASDAGV
SLGAAAAAHYRLTGSTRRRQVPYDDYLGHPYTQDDHDAAIGLVGDRVVTEPLELEAVA
DRIAAGDVVGWFQGGSEFGPRALGHRSVLADARSRDTWDFINAHIKFREDFRPLAPIV
PEEVAAEYFDLDEPSPHMLRVVPVREKYREQLAAVTHVDGTARVQTVSRDMNPRIHE
LLHLVGARTGFPVLVNTSLNRRGEPMIETPGQALDMLLGTRLSAMVLGDRVVRRAPEN
EAPLTLRSRLVLAPGVRLRWEQDCEGSRSWITGGAEPAGLELPRWAFDALSRAEPGR
ALGDYLPDCLVSAQGGTDTALAFLTALRARCLLVVTREAVHD

*DNA sequence*

GTGGACCCCGTCCATCTCCCGTCCGTCCGGCCTGGCCACGGCAGGTACCGCATG
ACACACGCACAGCAGCCCGCCTCGCCGCCGTGGTCCGTACTGAGCCCGGAACCC
GACCGCGGCGCCGGGCTCGGGGAAGGCGTACGTGTCCTGGTCGTCTCGGACGG
CACTCCCGCGCCCGACGGCGACGGGATGCCCGACGCGCACACGGAGCGGGTCG
CCGGACTGTGCCGGCAGCTGGCCGGCCGGGCGGAGATCGGTTACGCCTCGGCC
GCACGGCTGAGGGCCCGCAGCCTCGCCGCCGACCTGCGTGCGATGGTCACGGC
GGACGGCGGCTGGGACATCGTGGCCATGCCGTGGTCGTCGCCCGAGCGTGACG
CGGAACGGCTCAAGGTGCGGCGGGCCTTCGAGGAGGTGGCCCGGGACCCGGAC
ATGCCGCTGTTCGTCGCCGCTGCCGGACACGACGGACCGGGACGGCTCAGGTTC
CCCGCCTCCTGCCCCTCCGTGCTCGCCGTCGGGGTCGGCTCCGACCACGGCGGA
CCGACGGCCTACTGCGGCACGGACGGCCTGGGCCGCAAGCCTCAGCTGCTGGTC
CCGGACGCCCGCTACGCCACCCGCTGCGCGGCCGGCCCCGCTCCCATGCGCGG
CACCTCCGCCGCCGTCGGCATCGTCGCCGGACTCGCCGCCGTACTCGCACAGCG
GCTGGGGAGCGACGGACGCCGGGTGCCCTCGCCGCTGCTGCGTGCCGCGCTGC

TGGCCTCCGCCGGGCCGGACGGAACGCTGGTGGGCACGCCGCTGCTGCGGCAG
CCGGGGACCGGTGCCGCAGAACCGTTCTGCTTCGAGCTGCCCGCCGCAGCCGCC
CGGCTGCGGCTGCGCCTGAGAACCGGCGCGGACACCGTTCGGATCGCAGCCGTC
GCCGCGGGCGCGGACGAGGTCGTCAGCGGGGACACCGCCCCGTTGTGGCTGCC
GCGCGCCGCCGAACTCTCCGTGACATGGGCGGCGGACGGCACGACCCGGCGGA
CCACCGGCGCCGGGTGGCTGGTGCTCGATGTCGCATCCGGCGCCGGGCGTGAC
ACGGTGATCACCCTGGAGTGCCCGGAACCGGTCACCGCCTATGTCGCCGTCGGC
GGCGCCGAGGCCGTGGAGCGGATCGACCCTCGCCTCGAGAAGACCGACGAGCC
CGGCGTGACCGCTCCGGCGTGGCGGGCCGGCACCGGCCGTCCCGTCGTCCTCG
GCCTGTCGGCCAGCCATGACGCCTCCGCCTGCCTGCTGCGCGACGGCAGCCCGC
AGGTCGCCGTGCAGCAGGAGCGGGTCACCCGCCGGAAACACGACGGAGTCGGC
CACCTCTCCTCCCGTGCGGCCGCCGACTACTGTCTCACCTCGGCCGGCCTGACC
GCCGACGACGTGGACGTCTTCGCCTTCAACGCACAACCGCTGCTGCCTGGTTACG
TGGGGCTGTCGGTGCCGTCCGCGGCGCAGTCGTTCGACCTGTTCGACCCCTTCG
ACGCGCGCACCGTCTACGTCTCGCACCACCTCGCCCACGCCTTCTCCGCGTTCTG
GGGCTCGCCCTTCGAGGAGGCCGTGGTCGTCGTCTGCGACGGCTCCGGAGGCTC
GGTGCTCGGCACGGACGACCTCCTGGTGGACGGTCCCGGACTGCGGGAGTACCT
CGCGCGGGGGCCCGAGGACCGGCTGCCGCGCATCCACGTCTTCTCCGTCTACCA
CTTCGACCGGGACGGCTACCGGCTGCTGTTCCGCGAGTGCGCCGACTCCTTCAAC
GTCCGCGTCGGCTCCTCCTCGATCGGCGAGACCTATGCGGCCGTGAGCCAGTAC
GTCTTCGGCGACTGGCAGGAGGGCAGCGGCAAGCTGATGGGCCTCGCCCCCTGG
GGGAAGGCCCGCAACGCCGGCCCCAGCCTGCTGGAGCCCGGCCCGCACGGCCT
GCCCCGCTTCCGCAGCGACTGGAAGGACACCTATCGCGCCCCCGGCTCCGGCGG
CCCGATGGACCACGCCGATCTCGCGGCCCGGATCCAGGCCGACCTCGAAGAGGC
ACTGGTGGCTCGCATGCGGTACGCCATGTCTCTGGTGCCCGGCTGCCGCAACCTC
GCCTACGCGGGCGGCGTCGCGCTCAACTCCGTGGCCAACGACCGCATCGCCCGC
GAGAGCGGAGCCGACCGCTTCTTCGTGTTCCCGGCGGCGAGCGACGCCGGGGTC
TCCCTGGGCGCCGCCGCAGCCGCGCACTACCGGCTCACCGGCTCGACCCGGCG
CCGGCAGGTCCCCTACGACGACTACCTGGGACACCCGTACACCCAGGACGACCA
CGACGCGGCCATCGGGCTGGTCGGCGACCGGGTGGTCACCGAGCCGCTGGAGC
TGGAGGCCGTGGCGGACCGGATCGCCGCGGGCGACGTCGTCGGCTGGTTCCAG
GGCGGATCGGAGTTCGGGCCACGCGCCCTGGGCCACCGGTCGGTCCTCGCCGA
CGCCAGGAGCCGCGACACCTGGGACTTCATCAACGCGCACATCAAGTTCCGGGA
GGACTTCCGCCCACTGGCGCCCATCGTGCCCGAGGAGGTCGCCGCCGAGTACTT
CGACCTCGACGAGCCCTCTCCGCACATGCTGCGGGTGGTCCCGGTCCGCGAGAA
GTACCGTGAGCAGCTCGCTGCCGTCACCCACGTCGACGGCACCGCCCGGGTGCA
GACCGTCTCGCGGGACATGAACCCGCGCATCCACGAACTGCTCCATCTCGTCGGC
GCACGCACCGGTTTCCCCGTCCTGGTGAACACCTCGCTCAACCGCCGTGGCGAG
CCCATGATCGAAACGCCCGGCCAGGCACTCGACATGCTGTTGGGCACGCGGCTG
AGCGCCATGGTGCTGGGCGACCGTGTGGTCCGTCGCGCCCCCGAGAACGAGGCA
CCGCTCACCCTGCGCTCCCGGCTCGTCCTCGCCCCCGGCGTGCGGCTGCGGTGG
GAACAGGACTGCGAGGGCAGCCGTTCGTGGATCACCGGCGGCGCCGAACCGGC
CGGCCTCGAGCTGCCGAGGTGGGCGTTCGACGCCCTGTCCCGGGCCGAGCCCG
GCCGTGCCCTCGGCGACTACCTGCCGGACTGCCTGGTCTCCGCACAGGGCGGCA
CCGACACGGCGCTGGCCTTCCTGACCGCGCTCCGTGCCCGCTGCCTGCTGGTGG
TCACCCGGGAGGCAGTGCATGACTGA

*DNA sequence (E. coli optimized)*
ATGGATCCAGTCCACCTACCCTCAGTTAGGCCGGGTCATGGCCGTTACCGCATGA
CCCACGCACAACAGCCGGCGTCTCCGCCTTGGTCAGTACTGAGCCCGGAGCCGG
ATAGAGGCGCAGGCTTGGGTGAGGGGGTACGCGTTCTGGTCGTGTCCGACGGTA
CTCCGGCTCCGGACGGCGATGGTATGCCGGACGCGCACACCGAACGGGTGGCTG
GCCTGTGTCGTCAACTGGCTGGCCGCGCGGAGATCGGTTATGCCAGCGCGGCTA
GGCTGCGTGCGCGCTCACTGGCGGCTGATCTGCGTGCTATGGTTACGGCTGACG
GCGGTTGGGACATCGTGGCTATGCCGTGGAGCTCTCCGGAACGGGACGCCGAAC
GGCTTAAGGTCCGTCGTGCATTCGAAGAGGTGGCGCGTGACCCGGATATGCCTCT
GTTTGTGGCGGCTGCCGGTCACGATGGTCCGGGCAGATTGCGCTTCCCGGCGTC
CTGTCCGTCTGTACTGGCGGTTGGTGTGGGCAGCGACCACGGCGGCCCGACCGC
GTACTGCGGTACCGACGGCCTTGGCCGCAAGCCGCAACTGCTGGTGCCGGATGC
ACGCTACGCTACCCGTTGCGCGGCGGGTCCGGCGCCGATGAGAGGTACATCTGC
GGCGGTCGGCATTGTGGCAGGGCTGGCGGCGGTGTTAGCTCAGCGTCTCGGTAG
CGATGGCCGTCGCGTCCCGTCACCGCTGCTACGTGCAGCTTTATTGGCGTCTGCT
GGTCCGGATGGTACGCTGGTGGGTACGCCGCTATTACGTCAACCGGGTACTGGTG
CCGCGGAACCTTTCTGCTTTGAGCTGCCAGCGGCGGCAGCTCGTCTGAGGCTTCG
GCTGCGCACCGGAGCCGACACCGTTCGTATCGCCGCTGTGGCAGCCGGCGCGGA
CGAAGTTGTTTCGGGCGACACCGCACCACTGTGGCTGCCGCGAGCTGCCGAACT
ATCTGTTACCTGGGCAGCCGACGGCACGACCCGCCGTACGACCGGTGCAGGCTG
GCTGGTCCTGGACGTGGCCTCTGGTGCCGGACGCGATACCGTTATTACCCTGGAG
TGCCCGGAACCGGTTACCGCATACGTGGCGGTGGGGGGCGCCGAAGCTGTGGAA
CGCATCGATCCGAGGCTGGAGAAAACCGACGAGCCGGGCGTTACAGCGCCAGCG
TGGCGTGCGGGCACCGGCCGCCCAGTTGTCTTGGGTTTGTCTGCCAGCCACGAC
GCGAGCGCATGTCTGCTGAGAGACGGCAGCCCGCAGGTCGCCGTACAACAAGAA
CGTGTGACACGTCGTAAACATGACGGCGTCGGGCACCTGTCCAGCCGCGCAGCG
GCAGACTACTGCCTGACCAGTGCTGGCCTGACCGCAGACGACGTGGATGTTTTCG
CCTTTAACGCACAACCGCTCCTGCCAGGCTATGTTGGTCTGTCCGTTCCATCGGCC
GCGCAAAGCTTCGACCTTTTCGACCCGTTTGACGCGCGCACGGTTTACGTGAGCC
ATCATCTTGCGCACGCATTCTCAGCATTCTGGGGTTCCCCGTTCGAGGAGGCAGTT
GTTGTCGTATGTGATGGCAGCGGTGGTAGCGTGCTGGGTACCGATGACCTGCTGG
TAGATGGTCCAGGCCTGCGTGAGTATCTGGCGCGTGGCCCGGAGGACCGTCTGC
CGAGAATCCACGTTTTTAGCGTTTATCATTTTGATCGTGACGGCTACAGGCTGCTG
TTCCGCGAATGCGCAGACAGCTTTAATGTTCGCGTGGGCAGCTCAAGCATTGGTG
AAACCTACGCGGCAGTGAGCCAGTATGTCTTCGGCGATTGGCAGGAGGGTTCTGG
CAAGCTGATGGGTCTGGCGCCGTGGGGTAAAGCACGCAACGCTGGTCCGTCTCT
GTTGGAGCCGGGCCCGCACGGTCTCCCGAGATTCCGTAGCGATTGGAAAGATAC
GTACCGTGCGCCGGGTAGCGGTGGTCCTATGGATCACGCCGACCTGGCGGCGCG
TATCCAGGCAGACCTAGAGGAGGCGCTGGTCGCACGTATGCGTTACGCCATGAGC
CTTGTTCCGGGGTGCCGTAACCTGGCCTACGCAGGCGGTGTTGCTTTGAACAGCG
TGGCTAATGATCGCATCGCACGTGAGAGCGGTGCGGACCGCTTCTTTGTTTTCCC
GGCGGCTAGCGACGCAGGTGTCTCCTTGGGCGCGGCTGCGGCGGCGCATTATCG
TCTGACCGGTAGCACCAGACGTCGTCAAGTTCCGTATGATGATTATCTGGGACACC
CGTATACCCAGGATGATCATGACGCTGCGATTGGTCTGGTGGGCGATCGCGTGGT
GACCGAACCGTTGGAGCTGGAGGCGGTTGCGGACCGTATTGCAGCCGGTGATGT

GGTTGGTTGGTTTCAGGGTGGATCCGAATTTGGTCCGCGCGCGTTGGGCCACCGT
AGCGTACTGGCGGACGCGCGTTCTCGTGACACTTGGGATTTTATCAACGCACACA
TTAAATTCCGTGAAGATTTTCGTCCGCTGGCCCCTATTGTCCCGGAAGAGGTTGCC
GCGGAATATTTCGATTTAGATGAACCGTCCCCGCATATGTTGAGAGTGGTTCCGGT
GCGTGAGAAGTACCGCGAGCAGCTCGCGGCTGTGACCCATGTTGATGGGACGGC
TCGTGTTCAGACCGTTAGTCGCGACATGAATCCGCGAATCCACGAACTGTTGCAC
CTGGTAGGTGCGCGTACCGGTTTTCCGGTGCTGGTCAACACCAGCCTGAATCGAC
GTGGTGAACCCATGATTGAAACCCCGGGTCAGGCTTTGGATATGCTGCTGGGTAC
GCGTCTGTCTGCCATGGTTTTAGGCGATCGTGTCGTGCGTCGTGCTCCAGAGAAC
GAAGCACCGTTGACCCTGCGTAGCCGACTGGTTCTCGCGCCGGGAGTACGTCTAC
GCTGGGAACAGGATTGCGAAGGTTCCCGTAGCTGGATCACCGGCGGTGCGGAAC
CGGCTGGCTTGGAGCTGCCGCGTTGGGCATTTGATGCTCTATCCAGAGCTGAGCC
GGGCAGGGCGTTGGGCGATTACTTGCCGGACTGTCTGGTTTCGGCTCAAGGTGG
CACTGACACCGCGTTAGCGTTCCTGACCGCACTGCGGGCGCGCTGCCTCTTGGTT
GTGACGCGCGAGGCAGTGCATGACTAA

**StdG (Multiple Cloning Site 1)**

*Amino acid sequence*

MTDLDDRMTDLDDTPRTHALAWLENTTDDATMLPCGTSHRSVPAADAVAKVWPLRH
RLGISRVTDLTPLDVVGIPVFSVTRPQARTGQITLCQGKGNTPVEALASALFEAVERHC
GSLARPTLTARPTELRAAGRTHLTAADLGLAEPVPDQPIEWIQGRDSRTGSPVLLPAAE
VIFPYTAPPGCLRPVRPSTTGLASGATLSEAVLHALYEVVERDATSRYLHGGPGRLVAL
DTVTGSAETELLRKYARAEIDTVVIDLTHTTVLPTFAVFLSDPGAAQAHLAVSGFGTHP
NAGVALRRALTEASQARATAIQGSREDLDRVAEVYRADPAEQRAAFLHRALQAKAEG
VTDMSTLPALPPRSTRDTLRHVAALLEEGGYDRVVHTDLTVPELGLPVAHVAVPGMVD
SVVEPLRSHRVAHQHVQA

*DNA sequence*

ATGACTGACCTGGACGACCGTATGACCGACCTGGACGACACGCCGCGCACGCAC
GCTCTGGCGTGGCTGGAGAACACGACCGACGACGCCACCATGCTGCCGTGCGGC
ACCTCGCACCGGTCGGTGCCTGCGGCCGATGCCGTCGCCAAGGTGTGGCCGCTG
CGCCACCGCCTCGGCATCAGCCGCGTCACGGACCTCACCCCGCTCGACGTCGTG
GGCATCCCCGTCTTCAGCGTGACCCGGCCTCAGGCCCGCACCGGGCAGATCACT
CTCTGCCAGGGAAAGGGCAACACACCTGTGGAAGCCCTCGCCTCCGCGCTGTTC
GAGGCGGTGGAGCGGCACTGCGGTTCCCTGGCCCGCCCCACACTGACGGCCCG
GCCGACCGAGCTGCGGGCCGCCGGCCGCACCCACCTGACCGCGGCGGACCTGG
GCCTGGCCGAGCCTGTGCCCGATCAGCCGATCGAGTGGATCCAGGGCCGGGACA
GCAGGACCGGCAGCCCCGTGCTGCTGCCCGCCGCCGAGGTGATCTTCCCGTACA
CCGCGCCGCCCGGCTGTCTGCGGCCGGTACGCCCGAGCACCACGGGCCTGGCC
TCCGGCGCCACGCTGAGCGAGGCGGTGCTGCATGCGCTGTACGAGGTCGTCGAA
CGCGACGCCACCTCGCGGTACCTGCACGGCGGGCCGGGCCGGCTGGTCGCCCT
CGACACGGTGACCGGATCCGCCGAGACGGAACTGCTGCGCAAGTACGCCAGGGC
CGAGATCGACACGGTCGTCATCGACCTCACCCACACCACTGTGCTGCCCACGTTC
GCGGTGTTCCTGTCCGACCCGGGTGCCGCCCAGGCCCACCTGGCGGTGTCCGGT
TTCGGTACACACCCCAACGCGGGCGTCGCCCTGCGCCGCGCCCTGACGGAGGCC
TCGCAGGCCCGTGCCACCGCCATCCAGGGCAGCCGGGAGGACCTCGACCGCGT

CGCCGAGGTCTACCGCGCCGACCCCGCCGAGCAGCGGGCTGCGTTCCTGCACCG
CGCCCTGCAGGCGAAGGCGGAAGGCGTCACCGACATGAGCACGCTCCCCGCCCT
GCCGCCGCGGTCGACACGCGACACACTGCGGCACGTGGCCGCGCTGCTGGAAG
AAGGCGGCTACGACCGGGTCGTGCACACCGATCTGACCGTCCCGGAACTCGGTC
TGCCCGTCGCCCATGTGGCGGTGCCCGGCATGGTCGACTCCGTCGTGGAACCCC
TCAGGAGCCATCGTGTGGCCCATCAGCACGTCCAGGCTTAG

*DNA sequence (E. coli optimized)*
ATGACTGACCTAGATGATAGGATGACAGATCTGGACGATACCCCGCGTACCCATG
CACTGGCATGGCTGGAGAACACCACCGATGATGCAACCATGCTGCCGTGTGGTAC
AAGCCATCGTTCGGTGCCGGCGGCCGACGCGGTCGCGAAGGTTTGGCCGCTGCG
CCATCGTCTGGGTATTAGCCGCGTGACCGATCTAACCCCGCTGGACGTTGTTGGT
ATCCCGGTTTTCAGCGTTACCCGTCCGCAGGCGCGCACGGGCCAAATTACCCTTT
GCCAGGGCAAAGGTAATACCCCGGTAGAGGCGCTGGCATCTGCTTTGTTTGAAGC
GGTCGAGCGTCACTGCGGTAGCCTGGCGCGTCCGACTTTGACTGCAAGACCAACA
GAACTGCGTGCGGCAGGTCGTACGCACTTGACGGCTGCGGATCTGGGCTTGGCG
GAGCCGGTGCCGGATCAACCGATTGAATGGATTCAAGGTCGCGACTCCAGGACG
GGTTCCCCAGTTCTGCTCCCAGCTGCGGAGGTCATCTTCCCGTATACCGCCCCTC
CTGGGTGCTTGCGCCCAGTGCGTCCGAGCACCACCGGACTGGCTTCTGGTGCCA
CCTTATCTGAAGCCGTGCTTCACGCCCTGTACGAAGTTGTGGAGAGAGATGCCAC
CAGCCGCTACCTGCATGGCGGTCCGGGCCGCCTGGTTGCGTTGGACACGGTTAC
CGGCTCCGCAGAGACAGAACTGTTGCGCAAGTACGCACGTGCCGAGATCGACAC
CGTGGTGATCGACCTGACCCACACCACCGTGCTGCCGACGTTTGCAGTGTTCCTG
AGCGACCCGGGTGCGGCGCAGGCGCACCTGGCTGTCAGCGGTTTCGGCACGCAT
CCGAACGCAGGCGTTGCTCTCCGTCGTGCGCTTACTGAGGCCTCGCAAGCTCGTG
CGACCGCGATCCAAGGTAGTCGGGAAGATCTCGACCGTGTTGCGGAAGTTTATCG
TGCGGACCCGGCGGAGCAGCGTGCGGCGTTTCTGCACCGCGCGCTGCAGGCGA
AAGCAGAGGGTGTCACCGACATGAGCACCCTGCCGGCGTTGCCACCGCGTTCCA
CTCGTGACACCCTGCGCCACGTGGCAGCGTTACTGGAGGAAGGCGGTTATGATC
GTGTGGTGCACACCGATTTGACCGTTCCGGAACTGGGCCTGCCGGTCGCGCACG
TTGCCGTCCCGGGCATGGTCGATAGCGTGGTTGAACCGCTGCGTAGCCATCGTGT
TGCTCACCAGCATGTACAGGCATAA

**<u>StdH (Multiple Cloning Site 2)</u>**
*Amino acid sequence*
MWPISTSRLRPIVFGAASVRPLDPALLAAVDLRPPVRRGDLLPLLDGDPGTVVLVDGLF
GGTMAVTPTECRQLLDAGWTVVGCSSMGALRAADLWPLGAIGIGDIFTLYRLGTLTSD
ADVAVALDPDDGHREVTVCTVHVRAVLAAAVEAGLIAPAQRARLARAAEGIHWSQRS
WTACRAVWSELGVAPTVLSSLLRLGREPRLHPKVRDAETCLRAVLAGDWLGDGATDT
RRCPACQKPLTAGMLFCSVCVSEAHETCL

*DNA sequence*
GTGTGGCCCATCAGCACGTCCAGGCTTAGGCCCATCGTCTTCGGCGCGGCGTCG
GTACGTCCCCTCGACCCCGCGCTGCTGGCAGCGGTCGACCTGCGGCCTCCGGTC
CGGCGCGGGGACCTGCTTCCGCTGCTGGACGGCGATCCCGGCACCGTGGTCCTC

GTCGACGGACTGTTCGGCGGCACGATGGCCGTCACCCCGACGGAGTGCCGTCAG
CTGCTCGACGCCGGCTGGACGGTGGTCGGCTGCTCCAGCATGGGCGCCCTGCGC
GCCGCCGACCTGTGGCCGCTCGGCGCGATCGGCATCGGTGACATCTTCACCCTG
TACCGGCTGGGCACGCTGACCTCCGACGCGGACGTCGCCGTCGCCCTGGACCCG
GACGACGGCCATCGGGAAGTGACCGTGTGCACGGTGCACGTCCGGGCCGTGCTC
GCGGCCGCCGTCGAGGCGGGTCTCATCGCCCCCGCCCAGCGGGCCCGGCTCGC
CCGTGCCGCCGAGGGCATCCACTGGTCCCAGCGGTCCTGGACGGCCTGCCGGG
CGGTGTGGAGCGAGCTGGGAGTCGCCCCCACGGTGCTCTCCTCACTTCTGCGGC
TCGGCAGGGAACCCCGGCTGCACCCCAAGGTGCGCGACGCCGAGACCTGCCTGC
GCGCTGTCCTGGCCGGCGACTGGCTCGGCGACGGGGCCACGGACACACGCCGC
TGTCCGGCCTGCCAGAAACCACTGACGGCAGGAATGCTGTTCTGTTCCGTCTGCG
TCTCCGAGGCGCACGAGACCTGCCTGTGA

*DNA sequence (E. coli optimized)*
ATGGTATGGCCCATAAGTACATCAAGACTAAGGCCGATTGTGTTCGGCGCTGCGA
GCGTTCGTCCGTTGGACCCGGCGTTGCTGGCTGCCGTTGATCTGCGCCCACCGG
TTCGCAGAGGCGACCTGCTGCCGCTCTTGGACGGCGACCCGGGTACTGTTGTCCT
GGTAGACGGCTTGTTTGGCGGTACCATGGCAGTCACCCCGACCGAATGTCGTCAG
CTGCTGGACGCGGGTTGGACGGTGGTTGGGTGCAGCTCCATGGGTGCTCTGCGC
GCTGCGGATCTGTGGCCTCTGGGTGCCATCGGCATCGGTGACATCTTTACCCTGT
ACCGCTTGGGTACGCTGACTAGCGATGCGGACGTTGCCGTGGCCCTGGATCCGG
ATGATGGTCATCGTGAGGTTACCGTCTGCACCGTTCACGTTCGGGCGGTGCTCGC
GGCTGCGGTGGAAGCAGGTCTGATTGCACCGGCACAACGTGCACGTTTAGCGCG
CGCAGCGGAAGGTATTCATTGGTCGCAACGTAGCTGGACCGCGTGCCGTGCCGT
GTGGTCTGAGCTGGGCGTGGCGCCAACCGTTTTAAGCTCTCTGCTGCGTTTGGGC
CGTGAGCCGCGTCTTCACCCGAAGGTGCGTGATGCAGAGACATGCCTGCGTGCG
GTCCTAGCTGGCGACTGGCTGGGCGACGGTGCTACCGATACCCGTCGCTGTCCG
GCGTGCCAGAAACCGTTGACCGCGGGTATGCTGTTCTGCTCCGTGTGTGTTAGCG
AGGCGCACGAAACGTGCCTGTAA
